# CipA mediates complement resistance of *Acinetobacter baumannii* by formation of a Factor I-dependent quadripartite assemblage

**DOI:** 10.1101/2022.02.17.480811

**Authors:** Julia I. Ries, Marie Heß, Noura Nouri, Thomas A. Wichelhaus, Stephan Göttig, Franco H. Falcone, Peter Kraiczy

## Abstract

Multidrug-resistant *Acinetobacter baumannii* is known to be one of the leading pathogens that cause severe nosocomial infections. To overcome eradication by the innate immune system during infection, *A. baumannii* developed a number of immune evasion strategies. Previously, we identified CipA as a plasminogen-binding and complement-inhibitory protein. Here we show that CipA strongly inhibits all three complement activation pathways and interacts with key complement components C3, C3b, C4b, C5, Factor B, Factor D, and in particular Factor I. CipA also targets function of the C5 convertase as cleavage of C5 was impaired. Systematic screening of CipA variants identified two separate binding sites for C3b and a Factor I-interacting domain located at the C-terminus. Structure predictions using AlphaFold2 and binding analyses employing CipA variants lacking Factor I-binding capability confirmed that the orientation of the C-terminal domain is essential for the interaction with Factor I. Hence, our analyses point to a novel, Factor I-dependent mechanisms of complement inactivation mediated by CipA of *A. baumannii*. Recrutiment of Factor I by CipA initiates the assembly of a quadripartite complex following binding of either Factor H or C4b-binding protein to degrade C3b and C4b, respectively. Loss of Factor I binding in a CipA-deficient strain, or a strain producing a CipA variant lacking Factor I-binding capability, correlated with a higher susceptibility to human serum, indicating that recruitment of Factor I enables *A. baumannii* to resist complement-mediated killing.

**Significance Statement:** The Gram-negative bacterium *Acinetobacter baumannii* causes severe nosocomial infections and has developed various immune evasion strategies to overcome complement. The mechanisms how *Acinetobacter baumannii* successfully circumvents complement-mediated bacteriolysis are still poorly understood. Here, we show that the plasminogen-binding and complement inhibitory protein CipA terminates all three complement activation pathways. We describe a novel mechanism by which CipA cleaves the key complement components C3b and C4b in the presence of Factor H and C4b-binding protein by formation of a novel Factor I-dependent quadripartite complex. CipA, which has recently been successfully used for vaccination approaches, might represent an attractive target for the development of novel therapeutic interventions to block disorders with excessive hyperinflammatory complement activation.

## Introduction

*Acinetobacter (A.) baumannii* is considered as an emerging opportunistic pathogen of clinical significance and known to be a major cause of hospital-acquired infections (1–3). Of particular global concern and urgent health threat is the emergence of carbapenem-resistant or even pandrug-resistant *A. baumannii* (CRAB) (4–6). In 2017, the World Health Organization (WHO) has prioritized CRAB as critical pathogen for which drug research and development are urgently needed (7). In addition to extensive antibiotic resistance, the capability of this pathogen to overcome innate immunity enable *A. baumannii* to successfully establish infection in the human host (8, 9).

Complement is a central pillar of innate immunity and plays an important part in the defense against invading microorganisms, for the crosstalk between with immune cells as well as for homeostasis (10, 11). Activation of the complement system is typically achieved by three canonical pathways: the classical, lectin, and alternative pathway (12, 13). Antibody-mediated activation of the classical pathway (CP) involves initial binding of the C1 complex, while recognition of specific carbohydrate signatures results in activation of the lectin pathway (LP). The spontaneous activation of the key component C3, binding to cell surfaces or binding to properdin triggers the alternative pathway (AP). Binding of activated C3b to Factor B (FB) causes a change in the conformation that renders FB more accessible to cleavage by factor D (FD), thereby generating the soluble form of the C3 convertase of the AP. All three pathways converge into the assembly of the C3 convertases C3bBb of the AP or C4b2b of the CP and LP, respectively. Subsequent proteolytic cleavage of C3 into C3b and C3a by the formed C3 convertases leads to opsonization and flagging of invading microorganisms for phagocytosis with activated C3b molecules. Subsequent binding of C3b leads to the generation of the C5 convertases (C3bBb3b of the AP or C4b2b3b of the CP and LP) and thereby alters their substrate specificity toward C5. Cleavage of C5 into C5b and C5a initiates the terminal pathway by sequential binding of complement components C6, C7, C8, and multiple copies of C9 resulting in formation of the membrane attack complex (MAC). Finally, the integration of the pore-forming complex destabilizes the microbial membrane leading to killing of the intruding pathogen (12, 13).

To keep the activation of the complement system in check, all three pathways are tightly controlled by distinct complement regulatory proteins (12, 13). The assembly of the C3 and C5 convertases is controlled by soluble complement regulators such as Factor H (FH) or C4b-binding protein (C4BP). In circulation, FH and C4b-binding protein (C4BP) impairs the generation of these convertases by acting as cofactors for serine protease Factor I (FI)-mediated degradation of C3b and C4b (12, 13). Moreover, FI in solution exhibits a very low proteolytic activity toward C3b and C4b (14).

Several proteins have been described to contribute to complement resistance of *A. baumannii*, including AbOmpA (15), penicillin-binding protein-7/8 (PBP-7/8) (16), serine protease PKF (17), elongation factor Tuf (18), phospholipase D (19), surface antigen protein SurA1 (20), trimeric autotransporter Ata (21) as well as CipA (22), all of which are considered to play important roles in pathogenesis (23). Previously, we identified CipA as a complement inhibitory and plasminogen-binding molecule enabling *A. baumannii* to cross endothelial monolayers and to degrade the key complement component C3b by conversion of plasminogen to active plasmin (22). Of relevance, we also demonstrated that CipA also directly inhibits the AP, irrespective of its plasminogen-binding activity, and thus contributes to complement resistance of *A. baumannii*.

In this study, we sought to elucidate the underlying molecular principles of complement resistance mediated by CipA. Functional analyses revealed that CipA inhibits complement at the C3 and C5 activation level and impairs the function of the C5 convertases. CipA variants lacking FI binding capability completely lose their complement inactivating capacity indicating that interaction of CipA with FI is crucial for complement inhibition. Here we also show that a CipA-deficient strain, or a strain producing a CipA variant lacking FI-binding capability displayed a significantly higher susceptibility of *A. baumannii* to complement-mediated killing. Our analyses revealed a novel mode of complement inactivation mediated by CipA which involves the formation of a FI-dependent quadripartite-ordered assemblage consisting of CipA-FI-C3b-FH and CipA-FI-C4b-C4BP, respectively.

## Results

### The C-terminal region of CipA is responsible for the inactivation of the alternative and classical pathway

Recently, we showed that CipA of *A. baumannii* inactivates complement at early activation steps but did not impact the activation of the terminal pathway (22), e.g. by inhibiting the assembly of the MAC as previously described for different bacterial proteins (24–26). To gain deeper insights into the molecular principles by which CipA terminates complement activation and to narrow down the complement-interacting region(s), a number of single, double, and multi-point mutants as well as diverse deletion CipA variants were generated for functional analyses (Fig. 1). The inhibitory potential of CipA variants on the alternative and the classical pathway was assessed by preincubation of the purified proteins with different amounts of normal human serum (NHS) before adding to prepared microtiter plates to initiate complement activation. Deletion of 10 C-terminal amino acids (CipA ΔE360-K369) was sufficient to achieve a complete loss of the complement inactivating capacity of CipA on the AP and CP (Fig. 1B and C). However, CipA deletion variants CipA ΔA352-K369 and CipA ΔE360-K369 significantly enhanced activation of the CP (Fig. 1C). Substitutions or deletions of single or multiple amino acids did not affect or marginally affected the inhibitory potential of CipA on both pathways (Fig. 1E and F). CipA ΔQ19-Q144 carrying a N-terminal deletion of 144 amino acids retained complement inhibitory activity on both the AP and CP (Fig. 1B and D). This indicates that the whole C-terminal region (residues E360 to K369) but neither single amino acid residues nor electrostatic forces play a prominent role in complement inactivation as previously shown for variant CipA K365A-K369A that lose plasminogen-binding property (22).

**Fig 1.**
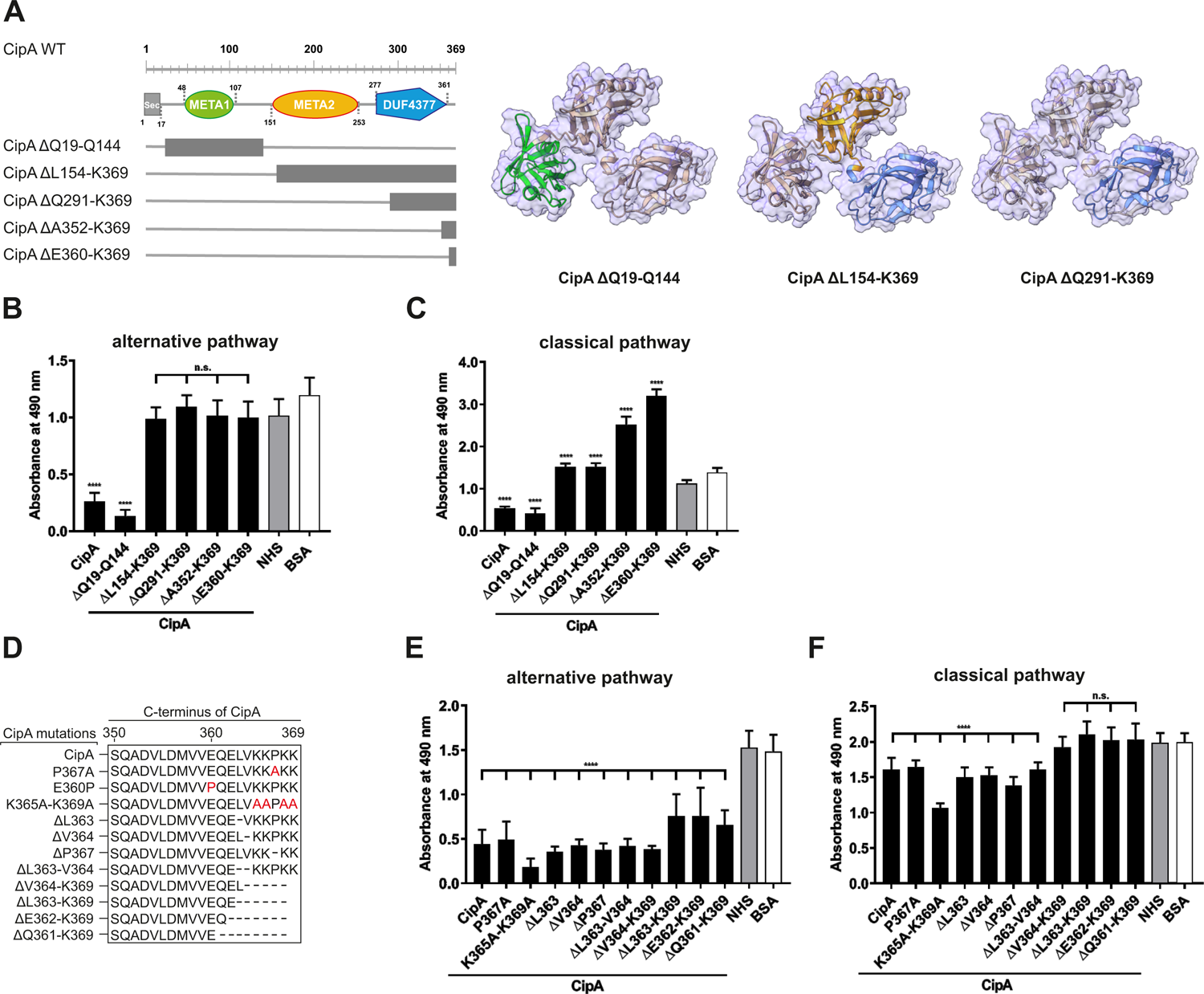
Assessment of the inhibitory capacity of CipA and CipA variants on activation of the AP and CP. Schematic representation of N- and C-terminal deletion CipA variants and structure predictions generating by using AlphaFold2.0 (A). Deletions are as follows: CipA ΔQ19-Q144 removes the whole META1 domain (green), CipA ΔL154-K369 removes the META2 (orange) and DUF4377 (blue) domains, while the CipA ΔQ291-K369 deletion removes most of the DUF4377 domain. Schematic representation of CipA variants carrying single and multiple deletions or substitutions (in red letters) (D). WiELISA were performed to assess the inhibitory capacity of CipA deletion variants on the AP (B and E) and CP (C and F). NHS pre-incubated with the purified CipA proteins or BSA (500 nM for AP and 2.5 µM for the CP) were added to microtiter plates immobilized with LPS (AP) or IgM (CP). Formation of the MAC was detected by using a monoclonal anti-C5b-9 antibody. Data represent means and standard deviation of at least three different experiments, each conducted in triplicate. ****, p ≤ 0.0001, n.s., no statistical significance, one-way ANOVA with post-hoc Bonferroni multiple comparison test (confidence interval = 95%).

To further determine the inhibitory potency of CipA on the AP, increasing concentrations of the protein were applied for ELISA. CipA showed strong dose-dependent, inhibitory properties on the AP with a calculated IC_50_ of 12.5 nM (Fig. S1A and S1B). This strong inhibition on the AP was also observed for the extracellular fibrinogen-binding protein (Efb) and its C-terminal fragment Efb-C of *Staphylococcus aureus* (Fig. S1C and S1D) (27).

### CipA of *A. baumannii* bound different complement components

The inhibitory capacity of CipA on the AP and CP suggests an interaction with various complement components. Employing ELISA, CipA bound C3, C3b, C4b and C5, but did not interact with complement regulator FH (Fig. 2). Although statistically significant, binding of C3c, FB, and FD to CipA was less pronounced. Further analyses of the CipA interacting complement components revealed a dose-dependent binding for C3b, C4b, and C5 (Fig. S2). As CipA appears to preferentially bind to C3b, we sought to narrow down the C3b-interacting region within CipA by comparing different CipA variants (Fig. S3). Unexpectedly, none of the modifications introduced in CipA showed a significant reduction in C3b binding. However, larger deletions at the N- and C-terminus displayed an inverse effect resulting in an increased C3b binding, in particular when variants CipA ΔQ19-Q144, CipA ΔL154-K369, and CipA ΔQ291-K369 were employed (Fig. S3B). In addition, minor changes of the C-terminus also did not affect C3b binding (Fig. S3D and S3E). These findings suggest that multiple regions within CipA are involved in binding of C3b.

**Fig 2.**
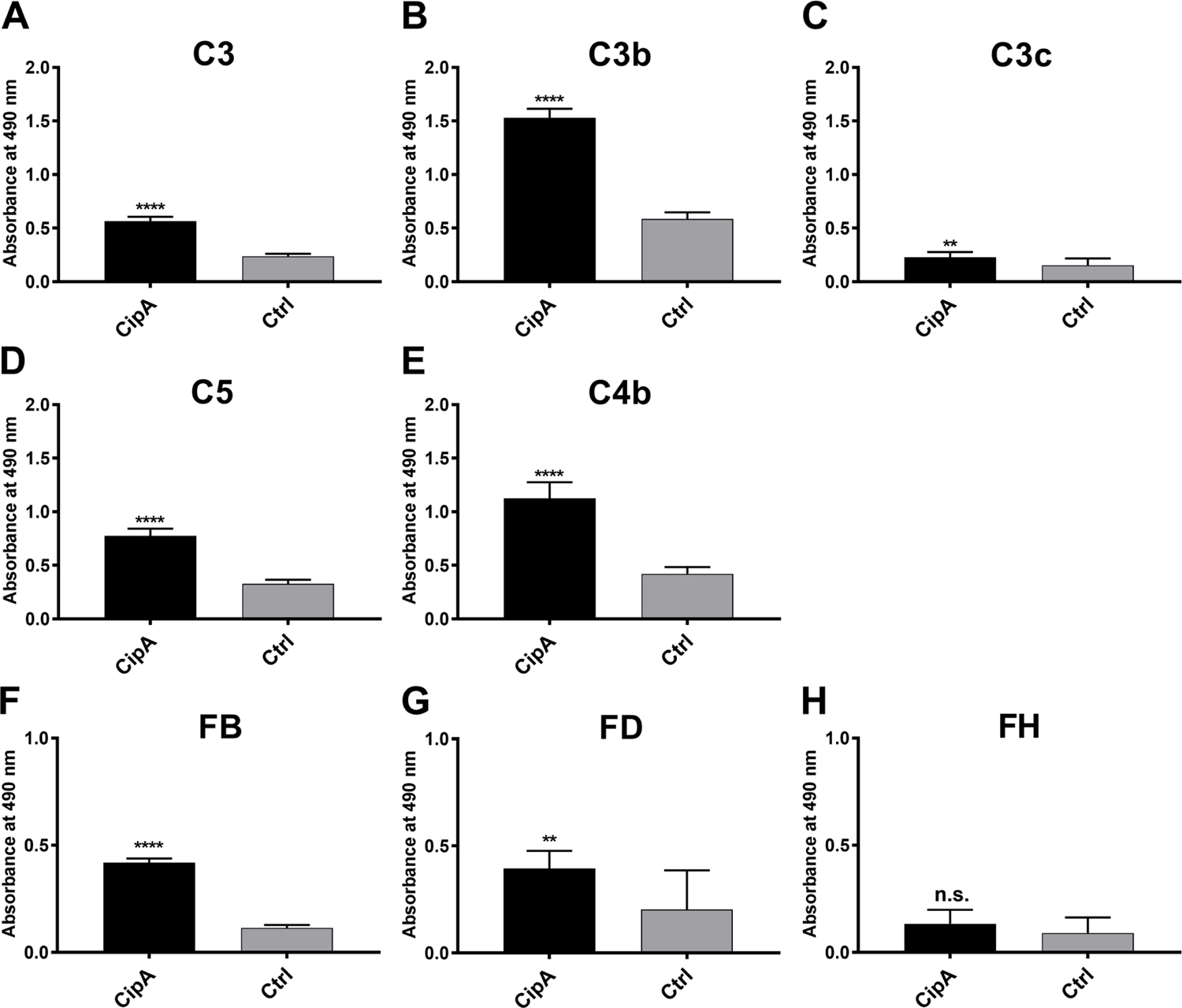
Binding of CipA to different complement components of the AP and CP. Protein binding of C3 (A), C3b (B), C3c (C), C5 (D), C4b (E), FB (F), FD (G), and FH (H), respectively, to CipA to was measured by ELISA. CipA (5 ng/µl) was immobilized and incubated with 5 ng/µl purified complement components and BSA or gelatine were used as negative controls (ctrl). Bound complement components were detected using specific antisera (1:1,000). To assess statistical significance, two-tailed, unpaired Student’s t-test was performed. Data represent means and/or standard deviation of at least three different experiments, each conducted in at least triplicate. **, p ≤ 0.05; ****, p ≤ 0.0001; n.s., no statistical significance.

### Interaction of CipA with the in vitro generated C3 convertase of the AP

Interaction of CipA with C3B and FB raises the possibility that CipA inhibits complement activation at the level of C3b by impeding the generation of the C3 convertase. In an initial attempt, a Ni-dependent C3bB proconvertase was assembled on microtiter plates as described (28, 29). A stable C3bB proconvertase could be generated *in vitro* upon detection of FB bound to C3b (Fig. S4A). In addition, dose-dependent binding of FB to C3b could be demonstrated by employing increasing concentrations of FB (Fig. S4B). Binding of FB to C3b was neither blocked when CipA was immobilized on microtiter plates prior adding C3b and FB (Fig. S4C) nor when CipA was added sequentially or in combination with FB (competitive) to C3b coated plates (Fig. S4D). Increasing concentrations of CipA of up to 2 µg/ml did not influence the generation of the surface-bound C3bB proconvertase (Fig. S4E).

To further investigate if CipA inhibits the C3bBb convertase formed in the fluid phase (29), the C3bB proconvertase was assembled by activating C3b-bound FB with FD in the presence and absence of CipA. Western blot analyses were investigated to detect cleavage of C3b generated by the assembled C3 convertase after 2 and 10 min and increased C3 concentrations (100 nM and 200 nM) (Fig. S4F-I). Collectively, the data of these analyses revealed that CipA did not have a strong impact on the generation of C3b (Fig. S4G and I) or targeted binding of FB to C3b (Fig. S4F and H).

### Interaction of CipA with the C5 convertases of the AP and CP

Having demonstrated binding of CipA to C5 (Fig. 2D), we investigated CipA-mediated impairment of the C5 convertases of the AP and CP by determining the generation of C5a employing ELISA. NHS was added to microtiter plates coated with either lipopolysaccharide (for the AP) or human IgM (for the CP) to induce the formation of the C5 convertases (30). Purified C5 preincubated in the absence or presence of CipA, variant CipA ΔE360-K369, and variant CipA E360P, respectively, were then added to the wells and the release of generated C5a in the fluid phase was detected. Generation of C5a was significantly impaired in the presence of the WT CipA, while variants CipA ΔE360-K369 and CipA E360P lacking complement-inhibitory activity did not impact cleavage of C5 (Fig. 3). These findings suggest that CipA facilitates complement inactivation at least at the level of C5b generation.

**Fig 3.**
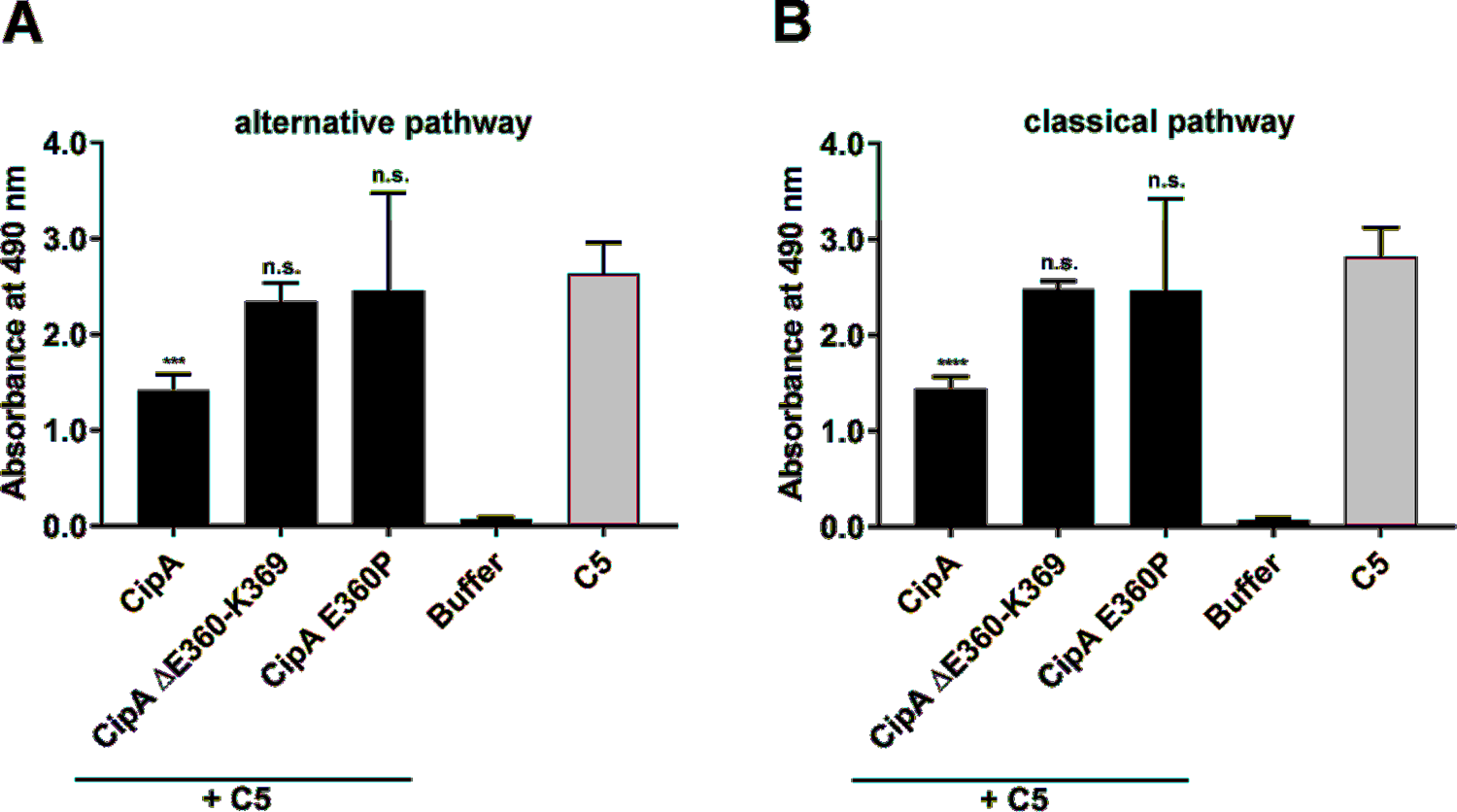
Impact of CipA on the enzymatic activity of the C5 convertases. The influence of CipA on the C5 convertase was analyzed by measuring the generation of C5a. The C5 convertases (AP and CP) were constituted on microtiter plates coated with LPS or IgM by adding NHS. Thereafter, reaction mixtures of C5 preincubated with or without CipA variants were added. The generation and release of C5a in the supernatant was then detected by C5a ELISA. Data represent means and standard deviation of at least three independent experiments, each conducted in duplicate. ***, p ≤ 0.0002; ****, p ≤ 0.0001, n.s., no statistical significance, one-way ANOVA with post-hoc Bonferroni multiple comparison test (confidence interval = 95%).

### CipA terminates complement activation by interaction with FI

The pronounced inhibitory capacity of CipA on the AP and the binding analyses indicate that an additional, yet unnoticed, molecular mechanism exists. Since we analyzed binding of the central components involved in AP and CP/LP activation, additional ELISA was conducted to investigate binding of CipA to FI. Thereby a strong affinity of CipA to FI with a dissociation constant of 3.0 nM (±0.28 nM) (Fig. 4A and B). By employing N- and C-terminal CipA variants, deletion of the C-terminal domain resulted in a completely abrogation of the FI interaction (Fig. 4C). Interestingly, deletion of single or double amino acids as well as substitutions of single or multiple amino acids did not influenced the interaction of CipA with FI (Fig. 4D). To ensure that all positions of potential relevance within the C-terminus had been considered, *in vitro* mutagenesis was conducted to replace amino acids at the remaining positions 360, 361, and 362, respectively, by alanine (Fig. 4E). Apparently, none of the newly generated CipA variants exhibited any change in FI binding compared to CipA. Interestingly, replacement of glutamic acid by proline but not by alanine at position 360 drastically abolished binding as observed for CipA ΔE360-K369 lacking the 10 terminal amino acids. This suggests that proline changed the orientation of the C-terminus in such a way that FI was unable to bind to CipA. CipA E360P significantly lost the capacity to inhibit activation of the AP, CP, and LP compared to CipA (Fig. S5), indicating that the C-terminus is relevant for FI binding and complement inactivation.

**Fig 4.**
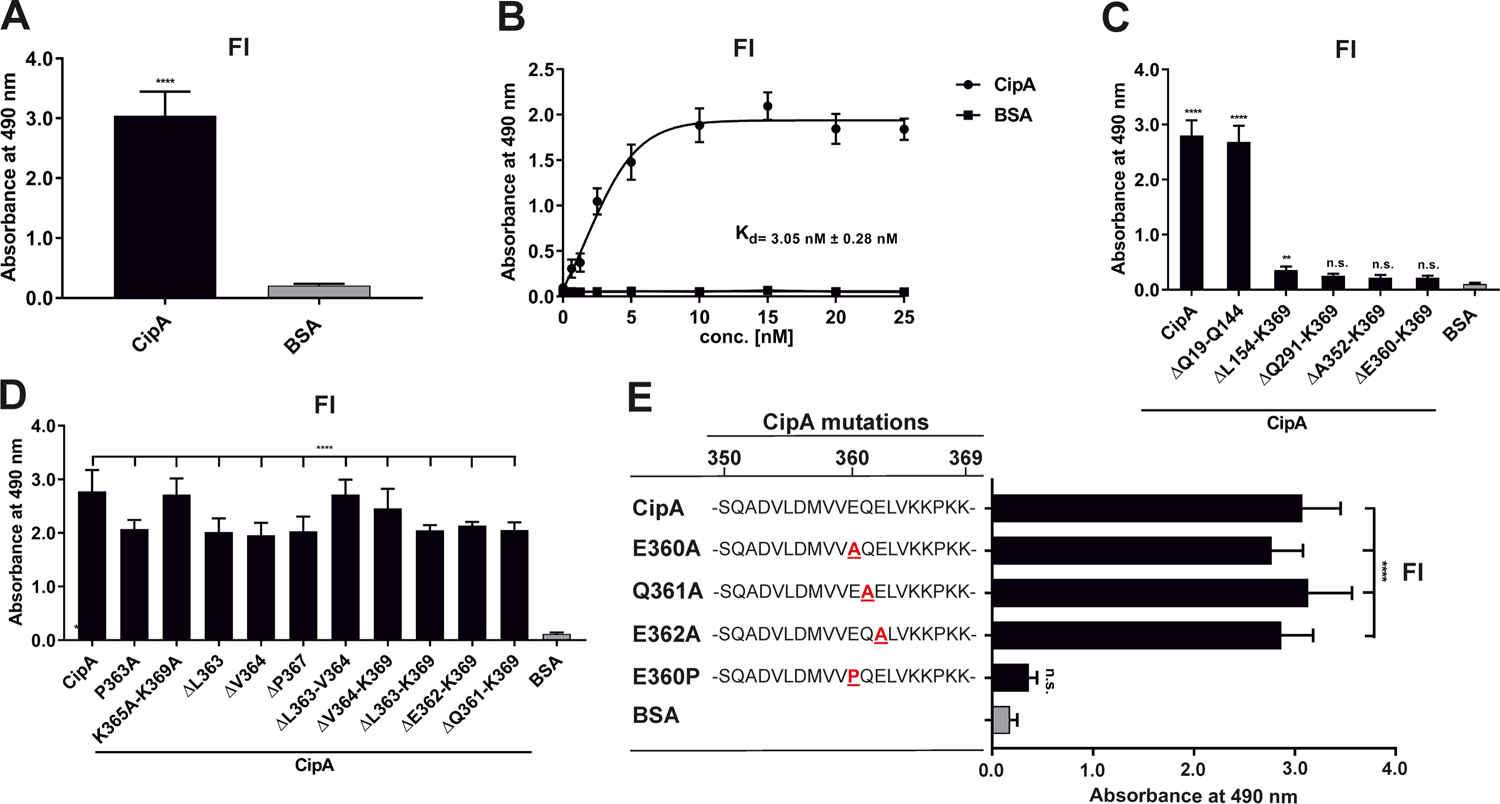
Interaction of CipA with FI. Protein binding of FI to CipA was measured by ELISA (A and B). CipA-coated wells (5 ng/µl) were incubated with 5 ng/µl purified FI and protein complexes were detected using an anti-FI antibody (1:1,000). To assess statistical significance, one-way ANOVA with post-hoc Bonferroni multiple comparison test (confidence interval = 95%) was performed. Data represent means and standard deviation of at least three different experiments, each conducted in at least triplicate. ****, p ≤ 0.0001. Dose-dependent binding of FI to CipA (B). CipA (5 ng/µl) immobilized was incubated with increasing concentrations (0 to 25 nM) of purified FI and dissociation constant was approximated via non-linear regression, using a one-site, specific binding model. Data represent means and standard deviation of at least three different experiments, each conducted in triplicate. Detection of the FI-interacting region within CipA employing diverse CipA variants (C to E). Microtiter plates coated with CipA and CipA variants (5 ng/µl) were incubated with FI (10 ng/µl) and binding was detected by an anti-Fi antibody (1:1,000). BSA was used as negative control in all assays. Data represent means and standard deviation of at least three different experiments, each conducted in triplicate. **, p ≤ 0.05, ****, p ≤ 0.0001, n.s., no statistical significance, one-way ANOVA with post-hoc Bonferroni multiple comparison test (confidence interval = 95%).

To further confirm the significance of the C-terminus in the interaction with complement, sequence and structural predictions were conducted (Fig. 5). SignalP5.0 (31) predicts a lipoprotein signal peptide (Sec/SPII) with a cleavage site between positions 17 and 18 (LMA-CQ) with a very high probability of 0.9991. The first 17 amino acids were therefore removed for structural predictions. AlphaFold2 (32) predicts a protein consisting of a short, unstructured N-terminal region, followed by three approximately equally sized domains with very high confidence (Fig. 5A and B). The three domains were identified by CDD/SPARKLE as two META domains (pfam03724) and one C-terminal DUF4377 domain (pfam14302; DUF, domain of unknown function). A strongly conserved DUF4377 is also found in other pathogenic bacteria associated with respiratory disease, such as *A. nosocomialis* (HCU39178; 99% sequence identity), *Klebsiella pneumoniae* (SSW87676.1; 100%), or bacteria related to non-respiratory diseases, e.g. Enterobacter asburiae (WP_194305131.1; 100%). META domains were first described in metacyclic stages of *Leishmania* (33), where the meta 1 gene is thought to be associated with virulence. The META1 domain (green) consists of 8 antiparallel beta-sheets flanked by two alpha-helices. A short linker connects it to the second META2 domain (orange), which has a similar arrangement of 8 anti-parallel beta sheets, but only one alpha-helix. This is connected by another short linker to the DUF4377 domain (blue), which consists entirely of beta-sheets. Glutamic acid at position 360 is located in the middle of the C-terminal beta-sheet strand of the DUF4377 domain (Fig. 5C). The introduction of a proline in position 360 disrupts the beta-sheet and dramatically alters the orientation of the last 10 C-terminal amino acids (Fig. 5D). A disruption of the protein fold by substitution of E360 with a proline is also predicted by Missense 3D (34). The pronounced effects on the structure of the C-terminal end of CipA are best appreciated in a short movie (Fig. S8).

**Fig 5.**
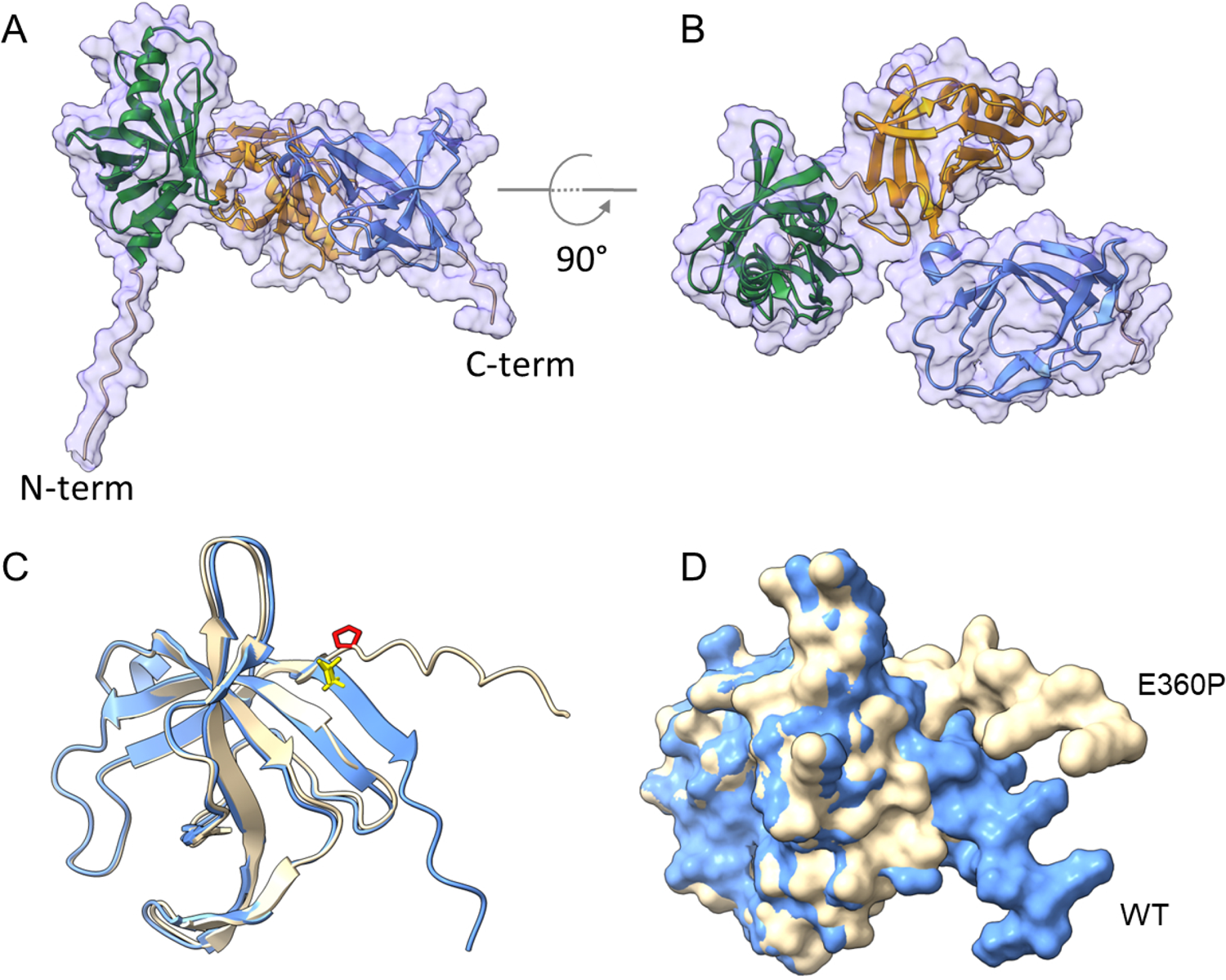
Structural prediction of CipA obtained with AlphaFold2. The mature CipA_18-369_protein has a short, 11 amino acid-long unstructured N-terminus (A), followed by three approximately equally sized domains (B), recognized as META domain 1 (AA 46-107), META domain 2 (AA 151-253) and a C-terminal domain of unknown function, DUF4377 (AA 277-361). The DUF4377 domain is made up entirely of beta-sheets, the last of which contains the E360 residue in WT CipA (blue, E360 highlighted in yellow in C) which was mutagenized to proline in the E360P mutant (beige, P360 highlighted in red in C). Introduction of this amino acid disrupts the final beta-sheet strands and results in reorientation of the last 9 C-terminal amino acids (D), away from the DUF4377 domain, but without any overt effect on the other parts of the molecule.

### CipA lacks intrinsic proteolytic activity but forms a quadripartite complex to terminate complement activation

To get a closer view into the molecular mechanism of complement inhibition mediated by CipA, we sought to exclude an intrinsic proteolytic activity of CipA as previously described for Pra1 of *Candida albicans* or SplB and ClpA of *Staphylococcus aureus* (35–38). CipA did not exhibit proteolytic activity on C3b and C4b when assayed under different experimental conditions (Fig. 6A to D). In addition, C3 was still unaffected while an additional band of app. 90 kDa appears when C4 was investigated (Fig. S6).

**Fig 6.**
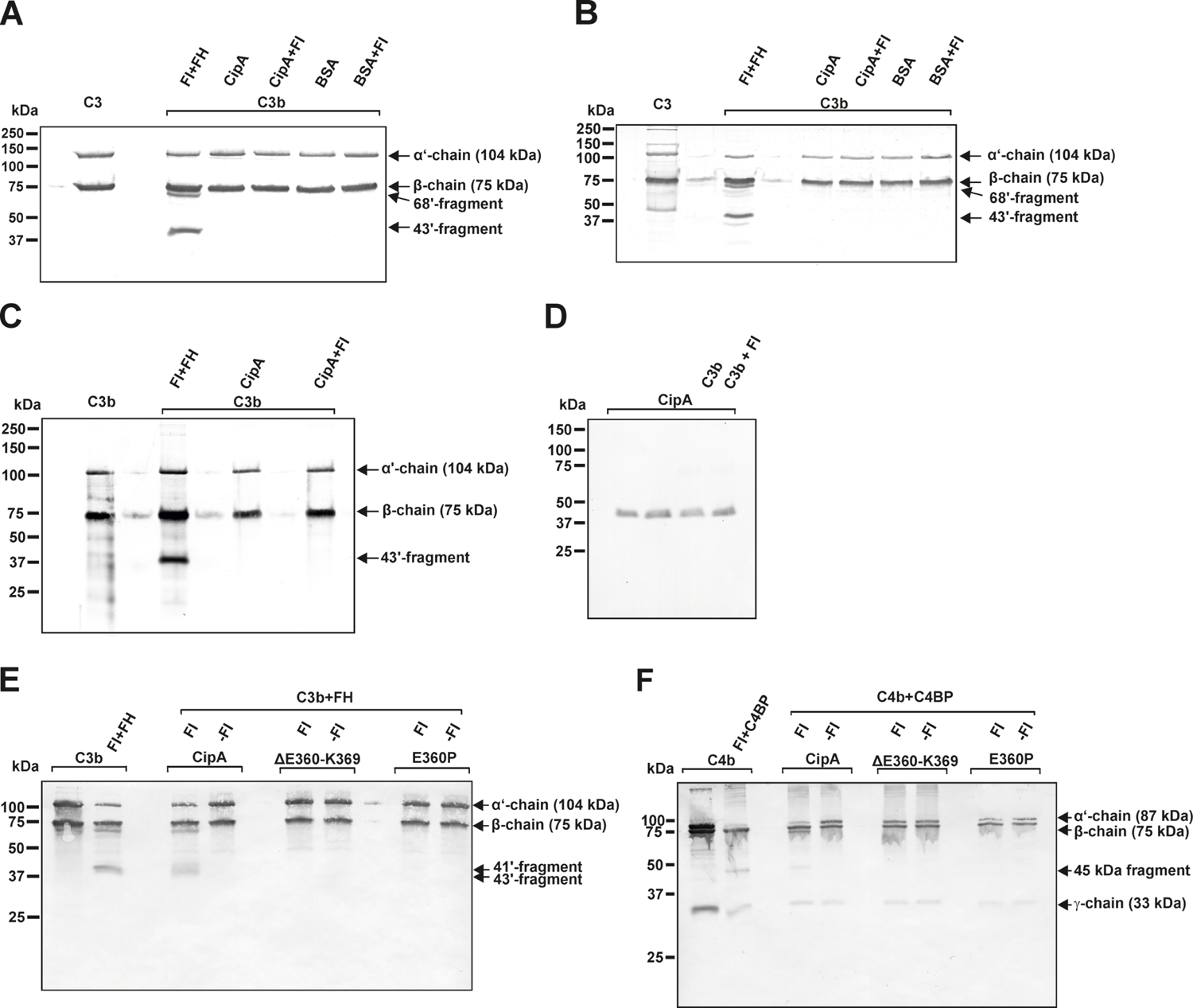
CipA lacks intrinsic proteolytic activity on C3b and C4b and formation of a quadripartite complex to inactivate C3b and C4b. Intrinsic proteolytic activity of CipA in FI-mediated inactivation of C3b was assessed by Western blotting (A to C). CipA or BSA coated wells were incubated with C3b or with C3b and FI for 1 h (A) and 6 h (B) at RT or for 6 h at 37°C (C). After termination, C3b degradation products were visualized by Western blotting employing a polyclonal anti-C3 antibody. Detection of CipA (D) in reaction mixtures shown in (C) by using an anti-CipA antibody (22). Formation of a quadripartite complex results in inactivation of C3b (E) and C4b (F). Microtiter plates coated with CipA, CipA ΔE360-K369 or CipA E360P (5 ng/µl) were incubated in the absence (-FI) or presence of FI (+FI). Thereafter, reaction mixtures containing C3b and FH or C4b and C4BP were added to the wells and were then subjected to SDS-PAGE and Western blotting. C3b and C4b degradation products were visualized by using an anti-C3 antibody and a mixture containing an anti-C4 and anti-C4d antibody. Control reactions containing C3b, FI, and FH as well as C4b, FI, and C4BP.

To further explore the nature of the complement inhibitory activity, CipA variants lacking Fi-binding capability were assayed with or without FI. Characteristic C3b and C4b cleavage products (41’ and 43’ fragments for C3b and 45 kDa fragment for C4b) could only be detected when FI bound to the immobilized WT CipA protein (Fig. 6E and F). By contrast, no degradation of C3b and C4b was observed in the absence of FI or when CipA variants lacking FI binding capability were investigated suggesting that cleavage occurs upon formation of a quadripartite complex consisting of CipA-FI-FH-C3b and CipA-FI-C4BP-C4b.

### Recruitment of serum-derived FI facilitates complement resistance of *A. baumannii*

To investigate the impact of FI on complement resistance of *A. baumannii*, the ΔcipA strain was complemented by either the WT *cipA* gene (*Ab ΔcipA::cipA*), a *cipA* gene with a deletion of 30 nucleotides at the C-terminus or a *cipA* gene with modifications at position 1078 (GAG → CCG; aa position 360: E → P) to produce variant CipA E360P using marker-less mutagenesis (22). Employing western blotting, *A. baumannii* 19606 WT as well as the complemented strains *Ab ΔcipA::cipA* and *Ab ΔcipA::cipA E360P* produced the respective CipA protein (Fig. 7A). As expected, no signal could be detected for *Ab ΔcipA*. In addition, a strong degradation pattern was observed for *Ab ΔcipA::cipA ΔE360-K369* suggesting that deletion of the C-terminus results in an enhanced susceptibility of CipA to proteolysis (Fig. S7).

**Fig 7.**
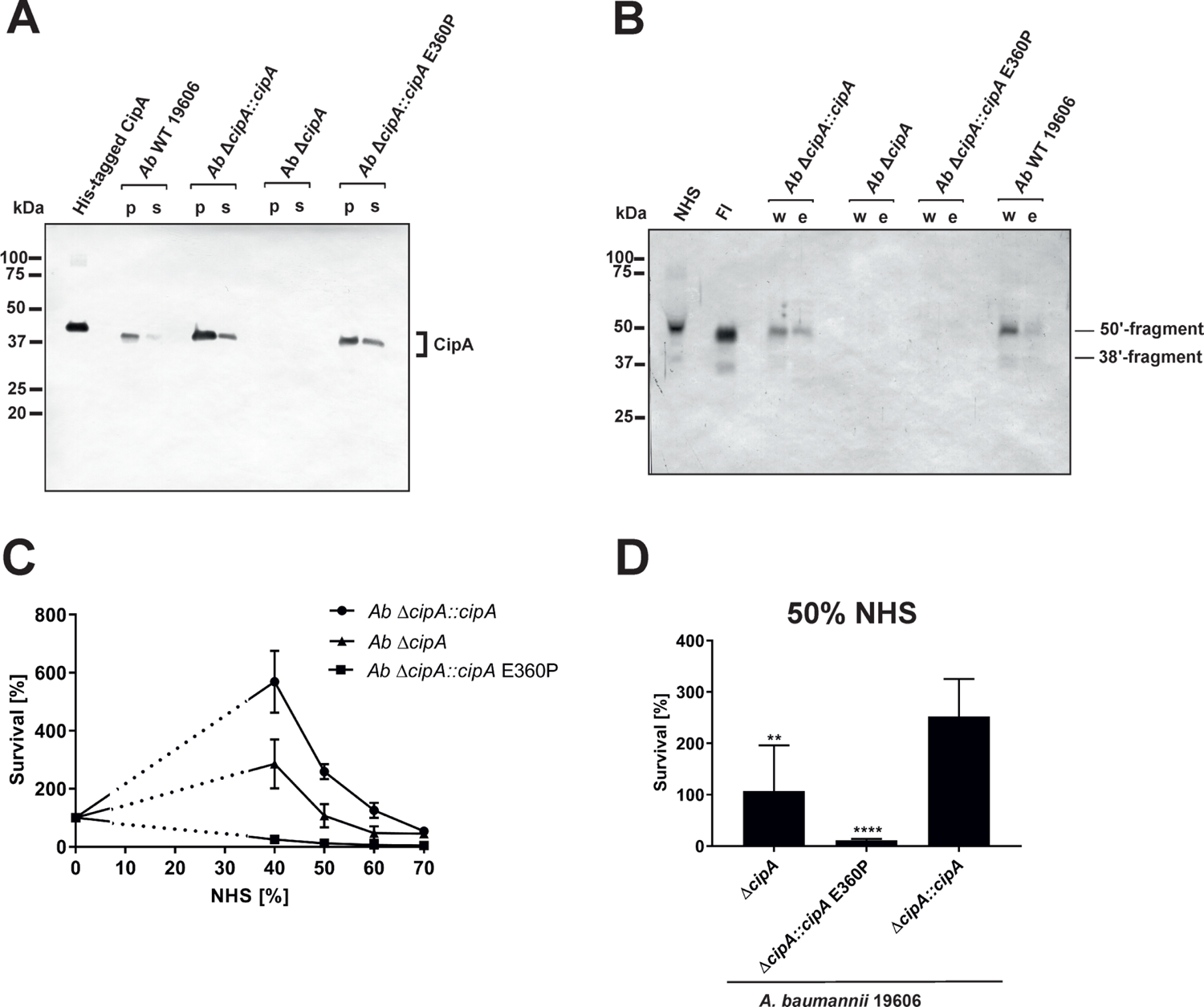
Recruitment of serum-derived FI to CipA mediates escape of *A. baumannii* from complement-mediated killing. CipA production in different *A. baumannii* strains was verified by Western blot analysis (A). CipA was detected in *A. baumannii* cell lysates (10 µg each) by using an anti-CipA antibody (1:100). Binding of FI to *A. baumannii* strains upon NHS incubation was assessed by Western blotting using an anti-FI antibody (1:1,000) (B). Survival of *A. baumannii* strains in human serum (C and D). 5 x 10 cells of *A. baumannii* ΔcipA::cipA (1), ΔcipA (▴), and ΔcipA::cipA E360P (▪) were incubated in increasing amounts of human serum for 2 h at 37 °C, serially diluted and cfu counts determined. Cfu counts were related to controls incubated in LB medium instead of serum, for which survival was set at 100%. Shown are results from at least three independent experiments (C). Survival of *A. baumannii* strains at 50% (D). **, p ≤ 0.002, ****, p ≤ 0.0001, n.s., no statistical significance, one-way ANOVA with post-hoc Bonferroni multiple comparison test (confidence interval = 95%).

Recruitment of serum-derived FI by *A. baumannii* was subsequently analyzed by incubating the three complemented strains with EDTA-NHS, and thereby surface-bound FI was detected by Western blotting (Fig. 7B). *A. baumannii* 19606 and *Ab ΔcipA::cipA* bound FI from human serum, whereas no binding could be observed for *Ab ΔcipA* and *Ab ΔcipA::cipA* E360P, suggesting that they are unable to bind FI.

Having demonstrated binding of FI by native *A. baumannii* cells, we sought to investigate serum susceptibility to explore whether interaction with FI contribute to complement resistance of this pathogen. Strains *Ab ΔcipA::cipA, Ab ΔcipA*, and *Ab ΔcipA::cipA* E360P were incubated with increasing NHS concentrations and survival was determined (Fig. 7C). Compared to the CipA WT producing strain *Ab ΔcipA::cipA*, strains lacking FI binding capability displayed an increased susceptibility to complement-mediated killing, in particular strain *Ab ΔcipA::cipA* E360P (Fig. 7D). Taken together, these data indicate that (i) CipA is the sole FI-binding protein of *A. baumannii*, (ii) recruitment of FI to the surface of *A. baumannii* occurs in the presence of all serum proteins, and (iii) CipA plays an important part in complement resistance of the pathogen.

## Discussion

*Acinetobacter baumannii* developed a number of versatile strategies to cope with the human immune system and thus, avoid recognition and eradication by complement as the first line of defense. Although numerous virulence factors and complement-affecting proteins have recently been described (15–23), the knowledge of the molecular mechanisms by which *A. baumannii* combat the effector function of complement is still in its infancy. Previously, we showed that CipA is able to inhibit complement activation by cleavage of the central component C3b following acquisition and activation of plasminogen (22). In this study, we expand our analyses and identify CipA as a multi-functional molecule of *A. baumannii* that interacts with diverse complement components including C3, C3b, C4b, C5, FB, and, in particular FI. CipA is capable of inhibiting all three activation pathways at the level of C3 and C5. Moreover, our analyses employing numerous CipA variants identified the C-terminus as the most relevant structural element involved in complement inhibition (Fig.1 and S1). CipA was also able to significantly inhibit the LP besides its property to inactivate the AP and CP (Fig. S3) indicating that inhibition occurs at the central hub of complement activation. Breakdown of the cascade by blocking the C3 convertases has also been reported for several immune evasion molecules secreted by *S. aureus* such as staphylococcal complement inhibitor (SCIN), extracellular fibrinogen-binding protein Efb, extracellular complement-binding protein Ecb, and the surface immunoglobulin-binding protein Sbi (27, 39–43). In contrast to the anti-complement activity of the staphylococcal immune evasion proteins, CipA does not inhibit the generation of the C3 convertases by binding to C3b and/or C3Bb or by formation of a tripartite complex consisting of CipA·C3b·FH (30, 40). Furthermore, Ecb is able to block C5 convertase activity by binding to C3b although C5 still binds to the formed C3 convertases (30). Here we showed that CipA binds to C3b and C5 (Fig. 2B and 2D) and impairs C5a generation (Fig. 3). Unlike Ecb, suppression of the enzymatic activity of the C5 convertases appears to follows a different mechanism of action as discussed below. CipA possess a strong inhibitory activity on the AP in comparison to Efb and Efb-C with a calculated IC_50_ of 14.5 nM, which is in the same range as reported for the staphylococcal proteins (Fig. S1C and D). Hence, the strong inactivation capacity makes CipA an attractive candidate in anti-inflammatory therapy. Future studies with appropriate infection models are necessary to prove whether this protein will serve as a suitable target for drug development.

So far, binding of serum FI has been described for *Prevotella intermedia* (44) and *S. aureus* (37, 38, 45, 46) whereby clumping factor A (ClfA) was identified as a cofactor for FI-mediated degradation of C3b (37, 38). Contrary to what we observed for the CipA-FI interaction, binding of FI to ClfA enhance FI-mediated cleavage of C3b into iC3b in the absence of complement regulator FH (Fig. 5). In addition, CipA did not display intrinsic proteolytic activities on C3/C3b and C4b as demonstrated for other microbial proteins such as Pra1 of *Candida albicans* (35), SplB and aureolysin of *S. aureus* (36, 47), and NalP of *Neisseria meningitidis* (48). These findings indicate that the molecular mechanism by which CipA promotes degradation of C3/C3b as well as C4b in the presence of FI is unknown.

Our structural predictions obtained with AlphaFold2 are particularly beneficial for interpreting the results obtained with the various CipA variants. Inhibition of the alternative and classical pathways do not appear to involve the META1 domain, as its complete removal does not show any effect (Fig. 1B and C). In contrast, removal of the META2 and DUF4377 domains or the DUF4377 domain alone results in complete ablation of CipA’s ability to inhibit the alternative pathway (Fig. 1B) and a partial reduction of the classical pathway inactivation capability (Fig. 1C). According to the AlphaFold2 structural prediction, the replacement of glutamic acid at position 360 with a proline results in a pronounced structural change in the C-terminal region of the DUF4377 domain, with complete loss of hydrogen bond pairing with the neighboring beta sheet. This alteration appears to have a strong impact on the ability of CipA to interact with FI (Fig. 4E) but did not affect interaction with C5 and FB (Fig. S9). Since substitutions with alanine at positions 360, 361 and 360, as well as deletions of single or multiple amino acids (Fig. 4D) had no effect on FI binding, the residues 360-369 are most likely not involved in direct FI binding. The dramatic rearrangement of the C-terminal portion in the E360P variant may instead disrupt other interactions between the DUF4377 domain and FI. The presence of a negative charge in position 360 also does not appear to play a role in the interaction with FI, as its substitution with a neutral alanine had no effect on binding (Fig. 4E). Of note, the structural rearrangement of the C-terminal beta sheet dramatically results in destabilization of this domain due to loss of several hydrogen bonds and thereby hamper FI accumulation (Fig. 5). Our data indicate that CipA assists FI in assembling of a quadripartite complex consisting of either CipA·FI·C3b·FH or CipA·FI·C3b·C4BP to mediated cleavage of C3b and C4b (Fig. 8). To our knowledge, this is the first description of a bacterial molecule involved in the formation of a proposed high-ordered complex by initial binding of FI and subsequent recruitment of C3b/C4b and their respective complement regulators to promote complement inactivation (Fig. 6 and 8). The C-terminal domain seems to stabilize the binding of free FI in the CipA·FI complex as a starting point for the subordinate enzymatic active assembly. In addition, the relevance of FI acquisition for complement resistance of *A. baumannii* was confirmed by the generation of CipA deficient strains producing either wild-type CipA or a CipA protein lacking FI-binding capability. The higher susceptibility of the FI-negative complemented strain (Fig. 7) strongly indicates that recruitment of FI enables *A. baumannii* to resist complement-mediated killing under more physiological conditions. These data provide novel insights into the underlying mechanisms of immune evasion of *A. baumannii*.

**Fig 8.**
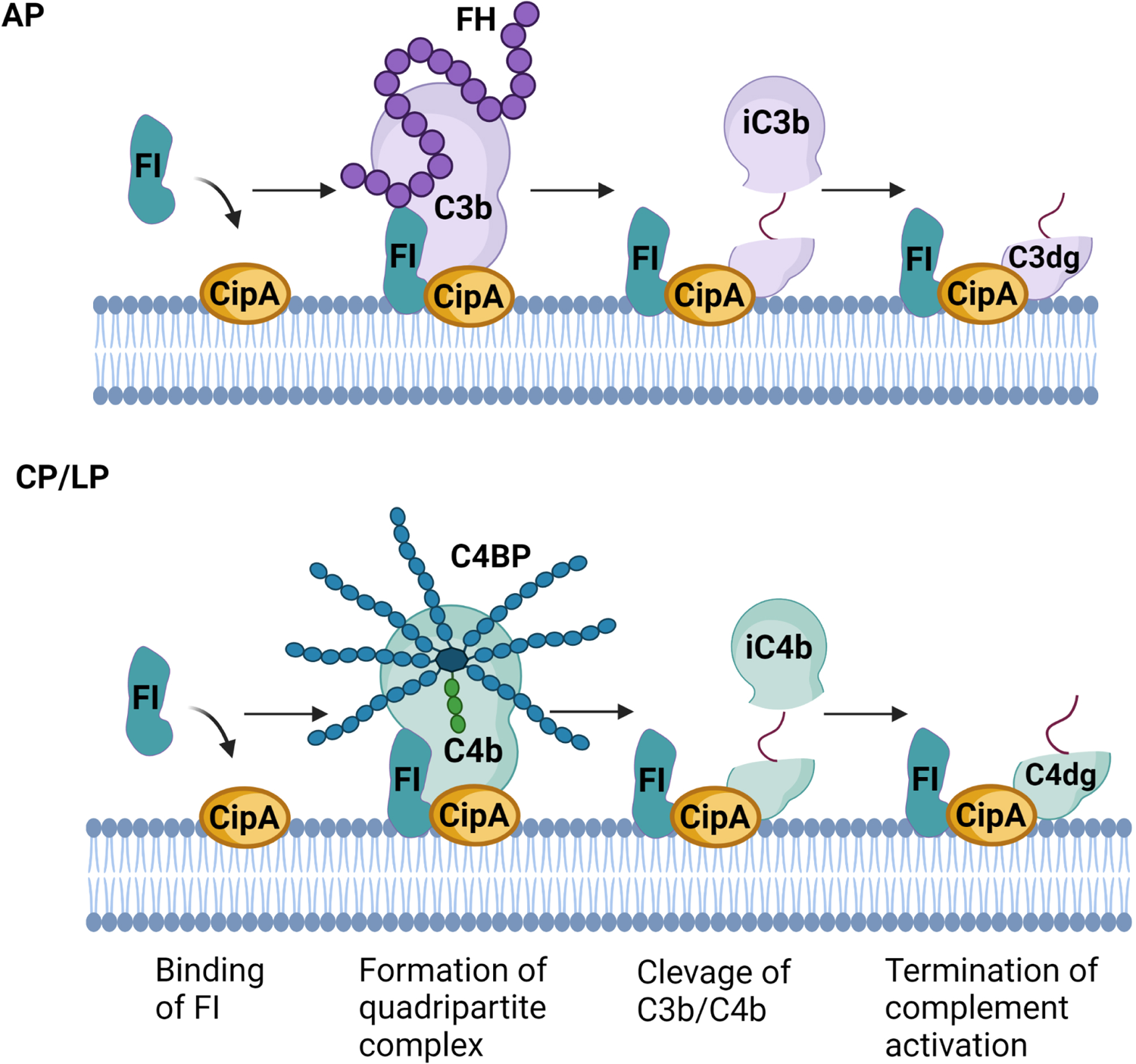
Molecular mechanism of complement inactivation mediated by CipA (hypothetical) Schematic representation of the molecular mechanism of complement inactivation by generating a quadripartite complex consisting of CipA, FI, C3b, and FH or CipA, FI, C4b, and C4BP. The figure was created with BioRender.com.

With the worldwide emergence of CRAB and the lack of appropriate antimicrobials on the horizon, novel therapeutic considerations including preventive measurements are mandatory. Passive and active immunizations with inactivated *A. baumannii* cells, cell fractions or outer membrane vesicles were able to protect mice from infection with multiple *A. baumannii* strains (49, 50). Formulations containing individual proteins as potential vaccines such OmpA, trimeric autotransporter Ata, serine protease PKF, 3-O-deacylase PagL, biofilm-associated protein Bap or poly-*N*-acetyl-β-(1–6)-glucosamine (PNAG) also provide protection of mice (51–56). More recently, first vaccination trials employing a vaccine containing CipA and the penicillin-binding protein PBP7/8 revealed that a combination of both proteins protects mice against infection with *A. baumannii* 19606 and induce strong cytokine responses towards IL-17 and IFN-γ (57). Due to its strong complement inactivating capacity, CipA is a promising candidate for the development of a suitable vaccine to prevent and treat serious infections caused by *A. baumannii*.

In conclusion, here we show that CipA (i) is a potent inhibitor of all three complement pathways (ii) acts as a key ligand for several complement components, and (iii) mediates formation of a quadripartite complex for FI-mediated cleavage of C3b and C4b in the presence of FH and C4BP. Acquisition of serum-derived FI appears to be an important mechanism of *A. baumannii* to evade complement-mediated bacteriolysis. The multi-factorial and strong anti-complement activity of CipA raises the possibility to target this molecule as a promising therapeutic drug for the treatment of *A. baumannii*.

## Materials and Methods

### Bacterial strains and culture conditions

*A. baumannii* strain ATCC 19606 (58), *A. baumannii* 19606 Δ*cipA* (22), *A. baumannii* Δ*cipA::cipA*, *A. baumannii* Δ*cipA::cipA* ΔE360-K369, and *A. baumannii* Δ*cipA::cipA* E360P were grown at 37 °C in lysogeny broth (LB) or on LB agar plates. *Staphylococcus aureus* USA300, *Escherichia coli* JM109, BL21 Star™ (DE3), and NEB 5-alpha, respectively, were grown at 37 °C in yeast tryptone broth or on LB agar plates.

### Proteins and antibodies

Purified complement proteins (C3, C3b, C4b, C5, FB, FD, FI, FH, C4BP) and the polyclonal anti-C5 antiserum were obtained from Complement Technology, Tyler, Texas, USA. Polyclonal antisera raised against complement C3, C4, and FH were from Merck (Darmstadt, Germany) and polyclonal antisera against FB and FI as well as the neoepitope-specific monoclonal antibody against C5b-9 were from Quidel, San Diego, USA. The monoclonal anti-FD antibody was from BioPorto Diagnostics A/S, Hellerup, Denmark. Anti-hexahistidine antibodies were from GE Healthcare, Chicago, Illinois, USA and Merck, respectively. Horseradish peroxidase (HRP)-conjugated immunoglobulins were obtained from Agilent Technologies Denmark, Glostrup, Denmark. The monoclonal anti-CipA antibody was described previously (22).

### Generation and purification of recombinant proteins

The hexahistidine-tagged CipA protein lacking the N-terminal signal sequence and encompassing amino acid residues 19-369 (CipA ΔQ19-Q144), C-terminally truncated CipA variants CipA ΔL154-K369, CipA ΔQ291-K369, and CipA ΔA352-K369, respectively, as well as variant CipA K365A-K369A were described previously (22). To introduce single, double or multiple amino acid substitutions or deletions into the C-terminal region of CipA, site-directed mutagenesis was conducted as described (59). Briefly, PCR was carried out for 18 cycles (95°C for 1 min, 55 °C for 1 min and 72 °C for 7 min) using 50 ng/µl pQE-CipA, 125 ng each of the oligonucleotides (S1 Table), and 4 U PCRBIO HiFi polymerase (PCR Biosystems). Following incubation with 10 U *DpnI* (New England Biolabs) to eliminate the remaining vector DNA, reactions were used to transform *E. coli* cells. Plasmid DNA obtained from selected clones was isolated and mutations at the desired positions were confirmed by DNA sequence analysis of both strands. Production of His-tagged proteins in either *E. coli* JM109 or BL21 Star™ (DE3) and subsequent purification by affinity chromatography using Ni-NTA agarose were performed as previously described (60). The purity of recombinant proteins was analyzed by 10% Tris/Tricine SDS-PAGE and silver staining techniques and the protein concentrations were determined by employing the Pierce™ BCA protein assay kit (Thermo Fisher Scientific, Rockford, IL, USA).

To generate a His-tagged Efb protein as an additional control displaying complement inhibitory activity on the C3 convertase (27, 61), the Efb-encoding gene of *S. aureus* USA300 was amplified by PCR using primers Efb_FP Bam and Efb_RP Sal (S1 Table). The amplified DNA fragment was then cloned into pQE-30 Xa (Qiagen, Hilden, Germany) yielding plasmid pQE-Efb. To generate a C-terminal fragment of Efb, plasmid pQE-Efb was used as template for PCR using primers pQE-RP in combination with Efb-C_FP_Bam. The resulting PCR product was cloned into pQE-30 Xa and the generated pQE-Efb-C plasmid was sequenced for verification.

### Enzyme-linked immunosorbent assay (ELISA)

To detect binding of complement components, Nunc MaxiSorp 96-well microtiter plates (Thermo Fisher Scientific) were coated with 100 µl of purified bacterial proteins (5 µg/ml) or BSA (5 µg/ml) in PBS at 4 °C overnight as described (62). Between every incubation step, wells were washed three times with PBS containing 0.05% (v/v) Tween 20 (PBS-T). After blocking with Blocking Buffer III BSA (AppliChem, Darmstadt, Germany) or with PBS containing 0.2% gelatine (w/v) (AppliChem), complement components (100 ng/ml) in PBS were added to the wells. Binding of complement components were then assessed by utilizing specific primary antibodies (dilution 1:1,000). Following incubation for 1 h at RT, HRP-conjugated anti-goat or anti-mouse IgG (dilution 1:1,000) were added and protein complexes were visualized using o-phenylenediamine (Merck). The absorbance was then measured at 490 nm employing PowerWave HT (Bio-Tek Instruments, Winooski, VT, USA). To determine dose-dependency and to calculate the dissociation constant, CipA was immobilized (5 µg/ml) and incubated with increasing amounts (0 to 50 nM) of C3b, C5, and FI, respectively. The antigen-antibody complexes were detected by using the appropriate antibodies as described above.

### Complement inactivation assays

To assess the inhibitory capacity of CipA and CipA variants on the classical (CP) and alternative (AP) pathway, an ELISA-based complement inactivation assay was conducted as described (63). Briefly, microtiter plates coated with either human IgM (30 ng/ml) (Merck, No I8260) for the CP, lipopolysaccharide from Salmonella enteridis (100 ng/ml) (Hycult Biotech Uden, The Netherlands) for the AP or mannan (1 µg/ml) (Merck) for the LP were incubated ON at 4 °C. After washing with TBS containing 0.05% (v/v) Triton X-100 (TBS-T), the plates were blocked with PBS-T containing 1% BSA for 1 h at RT. Between every incubation step, wells were washed three times with TBS-T. NHS (15% for the AP, 1% for the CP and 2% for the LP) was pre-incubated for 15 min at 37 °C with purified proteins (500 nM for AP and 2.5 µM for the CP and LP, respectively) and reactions were added to initiate complement activation. Formation of the MAC was detected by using a monoclonal anti-C5b-9 antibody (1:500) (Quidel, San Diego, CA, USA) followed by HRP-conjugated anti-mouse IgG (1:1,000). The reaction complexes were developed as described above.

### Generation of a solid-phase C3bB proconvertase

An ELISA-based approach was conducted to generate a Ni-dependent C3bB proconvertase as described (28, 29). Microtiter plates were coated with 3 µg/ml C3b in PBS overnight at 4°C, blocked with TBS-T containing 1% BSA, and incubated for 1 h at 37 °C with 800 ng/ml FB dissolved in phosphate buffer supplemented with 2 mM NiCl_2_, 4% BSA and 0.1% Tween 20. For initial experiments increasing concentrations of FB (0, 10, 20, 40, 80, 100, 200, 400, and 800 ng/ml) was employed. Binding of FB to C3b was detected using a polyclonal anti-FB antibody followed by a HRP-conjugated anti-goat IgG (1:1000). The reactions were developed as described above.

### Interaction of CipA with the C3bB proconvertase

The inhibitory capacity of CipA on the formation of the C3bB proconvertase was determined by using ELISA. Microtiter plates were coated overnight at 4 °C with CipA (5 ng/µl) in PBS and C3b (3 ng/µl) for control purposes respectively. After washing, wells were blocked with TBS-T containing 1% BSA and incubated for 1 h at 37 °C. After washing, C3b (3 ng/µl) dissolved in phosphate buffer supplemented with 2 mM NiCl_2_, 4% BSA and 0.1% Tween 20 was added and incubated for 1 h at 37 °C. Following washing, FB (8 ng/µl) resuspended in phosphate buffer supplemented with 2 mM NiCl_2_, 4% BSA and 0.1% Tween 20 were added and microtiter plates were incubated for 2 h at 37 °C. Binding of FB was detected using a polyclonal anti-FB antibody followed by a HRP-conjugated anti-goat IgG.

### Binding of CipA to the fluid-phase C3 convertase of the alternative pathway

To assess the ability of CipA to inhibit the assembly of the C3 convertase of the alternative pathway, the C3Bb complex was initially formed in solution as described (29). Briefly, C3b (200 nM), FB (100 nM), and FD (50 nM) were incubated in HBS-N (10 mM HEPES, 150 mM NaCl, 2 mM MgCl_2_, pH 7.4) for 2 min at RT. The reaction was then terminated by adding 5 mM EDTA and purified C3 was added at two different concentrations (100 nM and 200 nM) in the absence or presence of 1 µM CipA. Following incubation of 20 min at RT, Tris/Tricine incubation buffer was added and the proteins were separated by 10% Tris/Tricine SDS-PAGE and transferred to nitrocellulose. C3, FB, and cleavage products thereof were then detected by using anti-C3 and and FB polyclonal antibodies.

The CipA-mediated inhibition on the formation of the fluid-phase C3b proconvertase was investigated by preincubation of CipA (1 µM) with C3b (200 nM) for 15 min at RT. After adding FB (100 nM) and FD (50 nM) the reactions were initially incubated for 2 min at RT, and the reaction mixtures were then terminated with 5 mM EDTA. In addition, C3b (200 nM), FB (100 nM), and FD (50 nM) were incubated in the presence of CipA (1 µM) for 0, 2, and 10 min at RT. The reactions were then terminated by adding Tris/Tricine incubation buffer following separation through 10% Tris/Tricine SDS-PAGE. Detection of C3 and FB was performed by Western blotting as described above.

### Inhibition of the C5 convertases activity by CipA

The impact of CipA on the enzymatic activity of the C5 convertases was assessed by ELISA. Microtiter plates were coated overnight at 4 °C with either human IgM (30 ng/ml) for the CP or LPS (100 ng/ml) for the AP. After washing three times with PBS-T, wells were blocked with PBS containing 4% BSA and 0,1% Tween 20. To generate the C5 convertases, human serum (1% for the CP and 15% for the AP) was added and incubated for 1 h at 37 °C as described (30). After washing, the wells were then incubated with NHS for 1 h at 37 °C. Purified C5 (100 ng) and CipA (460 ng) as well as variants CipA ΔE360-K369 and CipA ΔE360P were pre-incubated in PBS for 30 min at 37 °C and then added to the wells for an additional incubation of 1 h at 37 °C. For control purposes, preincubated C5 was also added to the wells in the absence of CipA. Cleavage of C5 and detection of released C5a by the formed C5 convertases was detected using the MicroVue C5a EIA (Quidel) and the absorbance was then measured at 490 nm.

### Determination of the intrinsic proteolytic activity of CipA

To determine intrinsic proteolytic activity of CipA in FI-mediated inactivation of C3b, microtiter plates were coated with either CipA (5 ng/µl or 30 ng/µl) or BSA (5 ng/µl) overnight at 4 °C according to Hair et al. (38). After washing with PBS containing 0.05% (v/v) Tween 20 (PBS-T), wells were blocked with PBS containing 0.2% gelatine (w/v). Thereafter, reaction mixtures with C3b (1 µg) and FI (500 ng) or in which FI was omitted were added and the microtiter plates were incubated for different times points at RT and 37 °C, respectively. The reactions were then terminated by adding Tris/Tricine incubation buffer and mixtures removed were subjected to SDS-PAGE. After transfer to nitrocellulose membranes, C3b degradation products were visualized by a polyclonal anti-C3 antibody. As a positive control, C3b (600 ng) were incubated with FI (300 ng) and FH (10 ng) for 1 h at 37 °C and the reaction was then subjected to SDS-PAGE and Western blotting.

In addition, microtiter plates coated with CipA, CipA ΔE360-K369 or CipA E360P (5 ng/µl each) were incubated for 1 h at RT in the absence or presence of FI (5 ng/µl). After washing, reaction mixtures containing C3b and FH or C4b and C4BP were added to the microtiter plates. After incubation for 1 h at 37 °C reaction mixtures were transferred to tubes and reactions were terminated by adding Tris/Tricine incubation buffer. The samples were subjected to 10% Tris/Tricine SDS-PAGE following Western blotting. Control reactions containing C3b, FI, and FH or C4b, FI, and C4BP were incubated were incubated for 1 h at 37°C and for 2 h at 37 °C, respectively. The reaction mixtures were terminated by adding Tris/Tricine incubation buffer and subjected to SDS-PAGE. After transfer to nitrocellulose membranes, C3b and C4b degradation products were visualized by using a polyclonal anti-C3 antibody (1:1,000) and a mixture of a polyclonal anti-C4 (1:1,000) and a monoclonal anti-C4d antibody (1:100).

### Generation of *A. baumannii* strains producing diverse CipA variants

For complementation of *A. baumannii* 19606 ΔcipA, different vectors were generated harboring the (i) WT cipA gene, (ii) cipA gene with a stop codon for production of a CipA protein lacking 10 amino acids at the C-terminus (CipA ΔE360-K369), (iii) cipA gene with a substitution at aa position 360 (CipA E360P). Initially, the CipA encoding gene containing up- and downstream regions was amplified from *A. baumannii* 19606 by using primers CipA up fwd PstI and CipA down rev NotI. Following digestion with PstI and NotI, the resulting DNA fragment was cloned into the PstI/NotI sites of pCR 2.1 TOPO vector generating pCR_nc5_CipA_nc3. This vector serves as template for a subsequent PCR amplification with primers CipA V359 FP and CipA V359 RP to introduce two stop codons at aa position 359 and 360 in cipA. The DNA fragment was then digested with the respective restriction endonucleases and also cloned into pCR 2.1 TOPO vector resulting in pCR_nc5_CipA-359_nc3. Afterwards, the inserted DNA fragments were re-amplified, digested, and re-cloned into pBIISK_sacBkanR (64) resulting in vectors pBIISK_nc5_CipA_nc3 and pBIISK_nc5_CipA-359_nc3, respectively. In addition, pBIISK_nc5_CipA_nc3 was also used for *in vitro* mutagenesis to introduce a single substitution at position 360 (E to P) applying primers CipA E360P_II FP and CipA E360P_II RP. The vectors generated were then used to transform *A. baumannii* 19606 ΔcipA by a markerless mutagenesis approach as previously described (64). Briefly, kanamycin-resistant complemented cells were counter-selected with 10% sucrose to allow integration of the inserts into the genome of *A. baumannii* 19606 ΔCipA by homologous recombination. Selected clones were subjected to PCR analyses using primers CipA seq Fwd and CipA seq Rev to ensure that the fragments were correctly integrated and no additional mutations have been introduced in the CipA encoding gene and the 5’ and 3’ flanking regions during the cloning procedure. Sequence verification of cipA mutants was done by whole genome sequencing (WGS) as previously described (65). Briefly, DNA was extracted from isolates and WGS was carried out using the Illumina® MiSeq platform generating 250 bp paired-end reads with coverage of >50. Assembly and scaffolding after quality trimming of the reads was conducted using SPAdes v3.15.0. Sequences were mapped against the reference sequence (CP045110.1) and cipA was analyzed using Geneious 11.1.52.

### Recruitment of serum-derived FI by native *A. baumannii* cells

Binding of FI to *A. baumannii* 19606 WT, ΔcipA (22), ΔcipA::cipA, ΔcipA::cipA ΔE360-K369 and ΔcipA::cipA E360P was assessed by a serum adsorption assay as described previously (66). Briefly, bacterial cells grown to an OD_600_ of 0.5 were sedimented and resuspended in PBS^++^. Cells (1 × 10^10^) were incubated in 750 µl NHS-EDTA for 1 h at 37 °C. After washing, proteins bound to the bacterial surface were eluted by using 0.1 M glycine-HCl, pH 2.0. After adding 1 M Tris-HCl (pH 9.0), the cell debris were sedimented and the supernatant were analyzed by SDS-PAGE and Western blotting by using a polyclonal anti-FI antibody as described above.

### Serum susceptibility testing of *A. baumannii* strains

Serum susceptibility of *A. baumannii* ΔcipA, ΔcipA::cipA, and *A. baumannii* ΔcipA::cipA E360P was assessed by suspending 2.5 x 10 cells in PBS-diluted NHS (40% to 70%) following incubation at 37 °C for 2 h. Bacteria were serially diluted in PBS, plated onto LB agar plates and colony-forming units (CFU) were determined on the next day. The CFU counts were related to controls which were incubated with LB medium instead of NHS. The survival of bacteria in LB medium was set at 100% to compare the serum susceptibility of the complemented strains.

### Structure prediction analyses

The protein sequence of CipA (NCBI reference sequence WP 000696035.1) was used for structural prediction. After removal of the N-terminal 17 amino acid signal sequence, the mature protein sequence was submitted on the AlphaFold 2 advanced interface (67) with the default settings using the optional “Refine structures with Amber-Relax” option. Results were exported to UCSF ChimeraX 1.2.5. (68), using the Matchmaker function for the overlay of the DUF4377 domain in the WT CipA and variant CipA E360.

## Supporting information

Cover page for supplemental information

## Statistical analyses

The data collected represent means from at least three independent experiments, and error bars indicate SD. For statistical analyses, one-way ANOVA with Bonferroni’s multiple comparison post-hoc test (95% confidence interval) or a two-tailed, unpaired t-test were conducted by applying GraphPad Prism version 7.

## Ethics statement

The study and respective documents were approved by the ethics committee at the University Hospital of Frankfurt (control number 492/13). All healthy blood donors provided written informed consent in accordance with the Declaration of Helsinki.

## Acknowledgments

We thank Arno Koenigs for critical reading of the manuscript.

