## Supplementary figure 1 for "CipA mediates complement resistance of *Acinetobacter baumannii* by formation of a Factor I-dependent quadripartite assemblage"

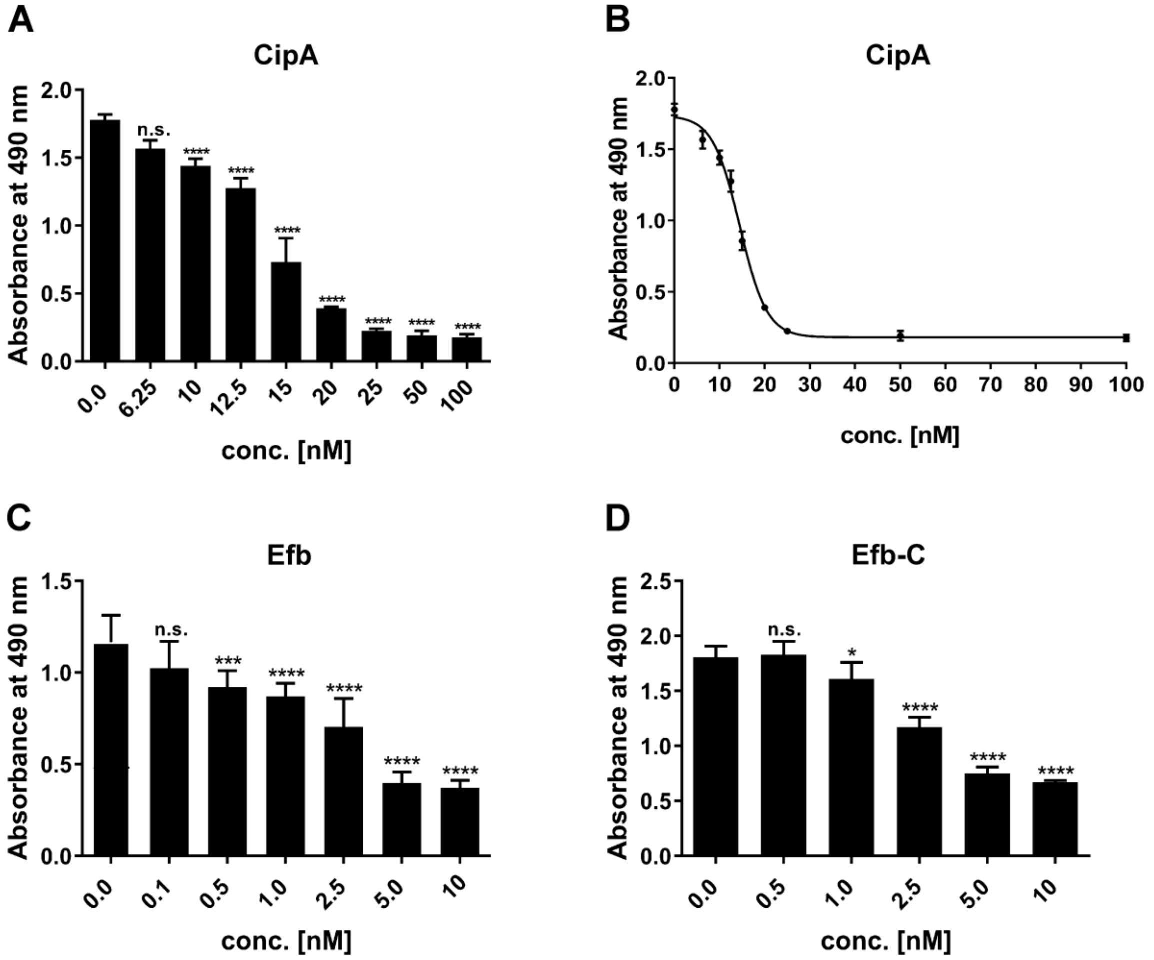


**Fig. S1. Dose-dependent inhibition of the AP.** Dose-dependent inhibition of the AP mediated by CipA **(A** and **B)**, Efb **(C)**, and Efb-C **(D)** was assessed by WiELISA. Microtiter plates immobilized with LPS were incubated with NHS prior incubated with increasing concentrations of bacterial proteins. The formation of the MAC was detected by using a monoclonal anti-C5b-9 antibody. Data represent means and standard deviation of at least three different experiments, each conducted in triplicate. *, $p \le 0.05$; ***, $p \le 0.0002$; ****, $p \le 0.0001$, n.s., no statistical significance, one-way ANOVA with post-hoc Bonferroni multiple comparison test (confidence interval = 95%). Binding curve and dissociation constant were approximated via non-linear regression, using a one-site, specific binding model using GraphPad Prism version 7.
