## Supplementary figure 2 for "CipA mediates complement resistance of *Acinetobacter baumannii* by formation of a Factor I-dependent quadripartite assemblage"

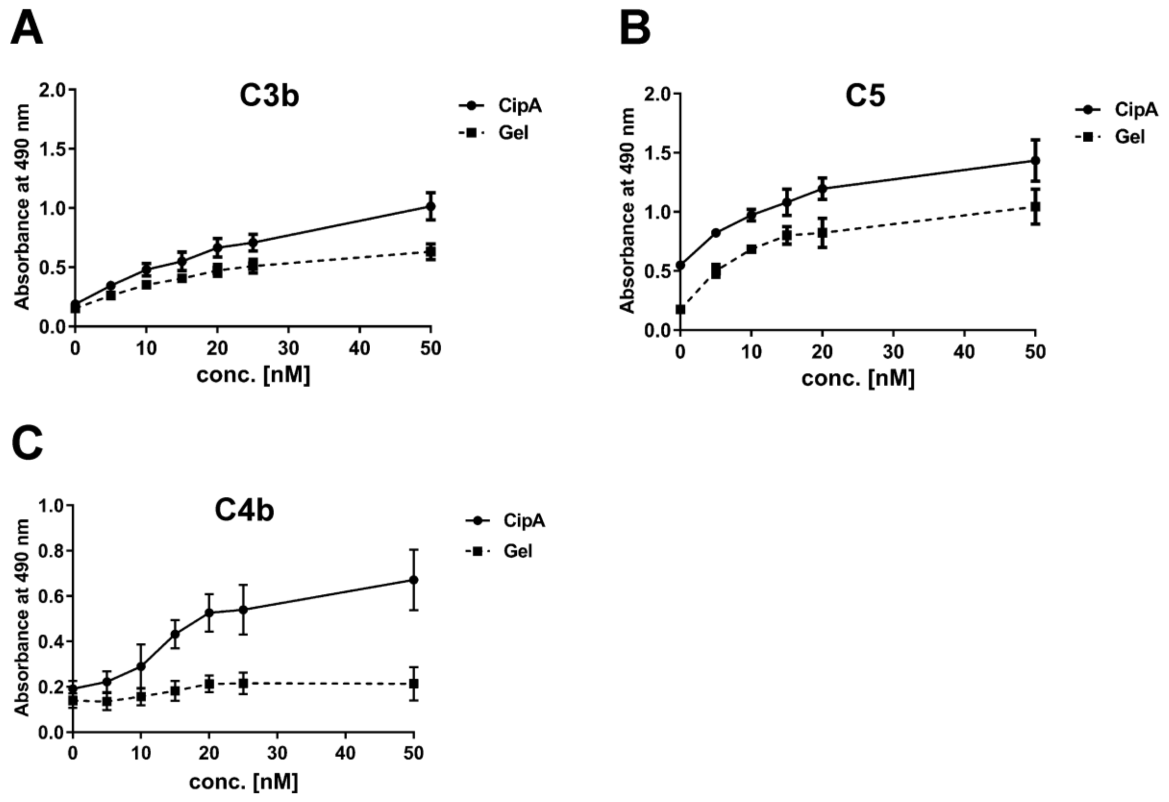


**Fig. S2 Dose-dependent binding of complement C3b, C5 and C4b to CipA.** Binding of C3b **(A)**, C5 **(B)**, and C4b **(C)** to CipA was assessed by ELISA. CipA (5 ng/µl) was immobilized and incubated with increasing concentrations (0 to 50 nM) of purified complement components. Data represent means and standard deviation of at least three different experiments, each conducted in triplicate. Gel, gelatine
