## Supplementary figure 3 for "CipA mediates complement resistance of *Acinetobacter baumannii* by formation of a Factor I-dependent quadripartite assemblage"

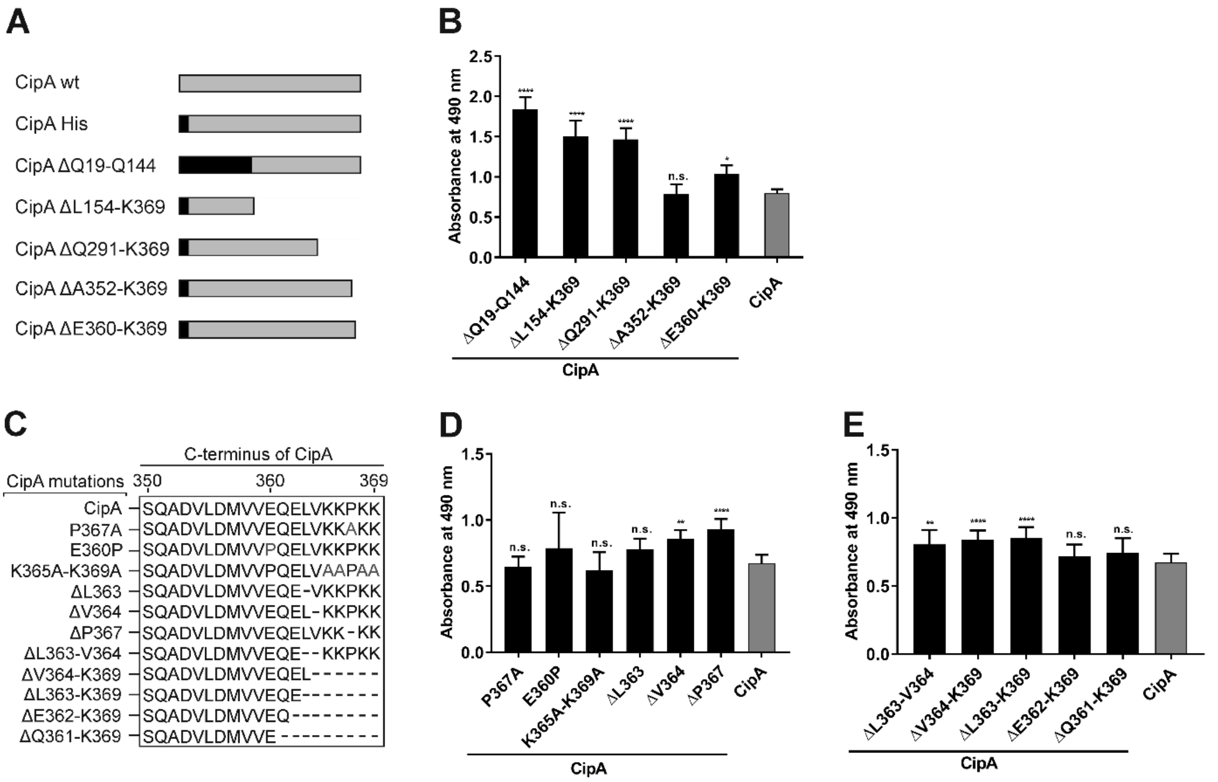


**Fig. S3 Binding of C3b to CipA variants.** Schematic representation of N- and C-terminal deletion CipA variants **(A)** the introduced deletions and substitutions within the C-terminus of CipA **(C)**. Protein binding of C3b to CipA variants was measured by ELISA **(B, D,** and **E)**. CipA (5 ng/µl) coated wells were incubated with purified C3b (5 ng/µl) and antigen-antibody complexes were detected using an anti-C3 antibody (1:1,000). Data represent means and standard deviation of at least three different experiments, each conducted in triplicate. *, p ≤ 0.05; **) p ≤ 0.005 ****, p ≤ 0.0001, n.s., no statistical significance, one-way ANOVA with post-hoc Bonferroni multiple comparison test (confidence interval = 95%).
