## Supplementary figure 4 for "CipA mediates complement resistance of *Acinetobacter baumannii* by formation of a Factor I-dependent quadripartite assemblage"

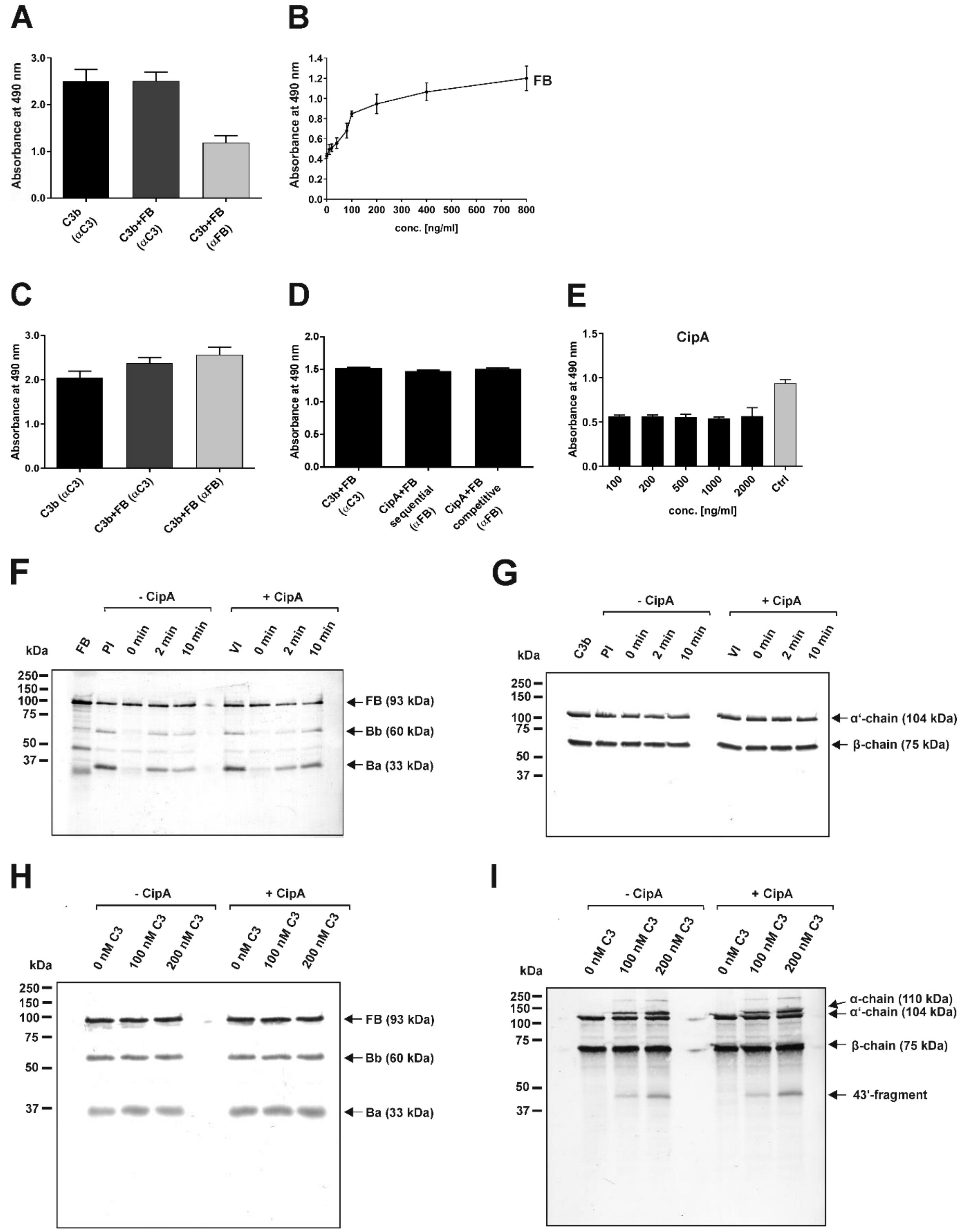


**Fig S4. Interaction of CipA with the C3B proconvertase and C3 convertase C3bBb of the AP.** Detection of a $Ni^{2+}$-dependent C3bB proconvertase by ELISA **(A** and **B)**. C3b-coated wells (3 ng/µl each) were incubated with either 500 ng **(A)** or increasing amounts of FB (0-800 ng/ml)

**(B)** in the presence of 2 mM $Ni^{2+}$ and analyzed by ELISA by employing an anti-C3 or anti-FB antibody. Data represent means and standard deviation of at least three different experiments, each conducted in at least triplicate. Impact of CipA on the formation of the C3bB proconvertase **(C** to **E)**. CipA-coated wells (5 µg/ml) were incubated with C3b (3 ng/µl) or with C3b and FB (3 ng/µl each) **(C)**. C3b-coated wells (3 ng/µl) were either incubated with CipA (5 ng/µl) and thereafter with FB (5 ng/µl) (sequential) or with CipA and FB (5 ng/µl each) (competitive) **(D)** or with increasing concentrations of CipA (0.1 to 2 ng/µl) **(E)**. Antigen-antibody complexes were detected by an anti-C3 or anti-FB antibody. Data represent means and standard deviation of at least three different experiments, each conducted in at least triplicate. Determination of the inhibitory capacity of CipA on the formation of the AP C3 convertase (**F** and **G**) and the cleavage activity of the formed C3 convertase **(H** and **I)** by employing Western blotting. The C3bB proconvertase was assembled by activating C3b-bound FB with FD in the presence (+CipA) or absence of CipA (-CipA) in the fluid phase. At different time points (0, 2, and 10 min) reactions were terminated by adding SDS sample buffer and subjected to SDS-PAGE. FB and C3b fragments were detected by Western blotting using appropriate antibodies. PI, preincubated. Cleavage of C3 by the assembled C3 convertase was detected by Western blotting. The formed C3 convertase was incubated with increasing concentrations of C3 (0, 100, and 200 nM) in the presence or absence of CipA and C3 cleavage products as well as FB fragments were detected by appropriate antibodies.
