## Supplementary figure 5 for "CipA mediates complement resistance of *Acinetobacter baumannii* by formation of a Factor I-dependent quadripartite assemblage"

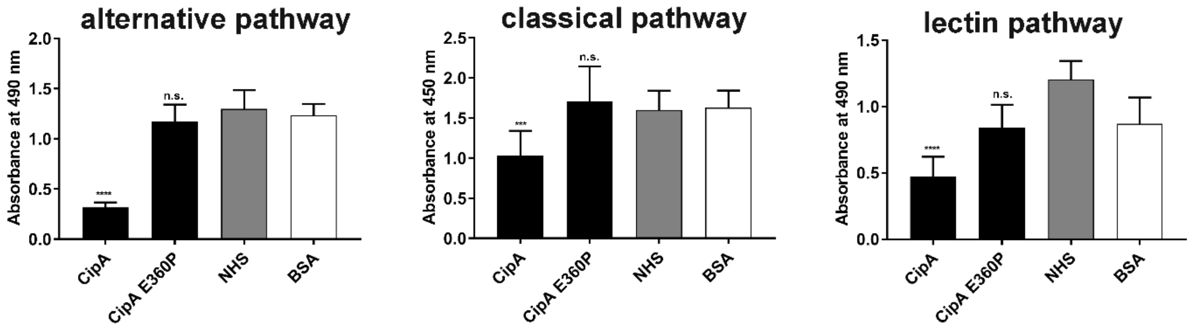


**Fig. S5 Comparative analyses of the inhibitory capacity of CipA and CipA E360P on the CP, and CP.** Inhibition of the three complement pathways by CipA and CipA E360P was assessed by WiELISA. Microtiter plates immobilized with LPS (AP), IgM (CP), and mannan (LP), respectively, were incubated with NHS prior incubated with purified proteins (500 nM for AP and 2.5 µM for the CP and LP, respectively). The formation of the MAC was detected by using a monoclonal anti-C5b-9 antibody. Data represent means and standard deviation of at least three different experiments, each conducted in triplicate. ***, $p \leq 0.0002$; ****, $p \leq 0.0001$, n.s., no statistical significance, one-way ANOVA with post-hoc Bonferroni multiple comparison test (confidence interval = 95%).
