## Supplementary figure 6 for "CipA mediates complement resistance of *Acinetobacter baumannii* by formation of a Factor I-dependent quadripartite assemblage"

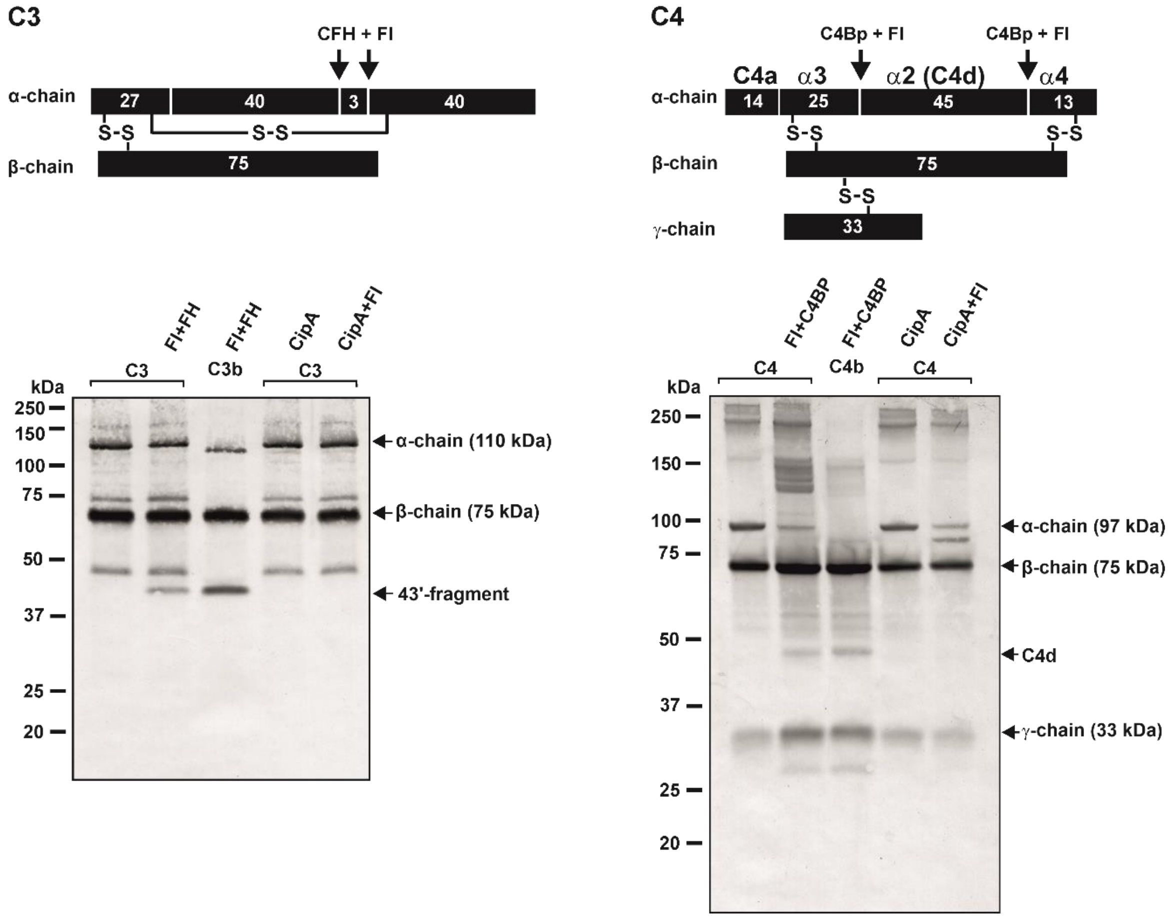


**Fig. S6 Determination of the intrinsic proteolytic activity of CipA on C3 and C4.** Intrinsic proteolytic activity of CipA was assessed by Western blotting. Schematic representation of C3 and the cleavage fragments generated by FH and FI (left panel) and C4 and the cleavage fragments generated by C4Bp and FI (right panel). CipA coated wells were incubated with C3 or C4, washed and reaction mixtures containing FH and FI or C4bp and FI were added. After incubation of 2 h at 37 °C, reactions were terminated, subjected to SDS-PAGE and, C3b and C4b degradation products were visualized by Western blotting using an anti-C3 antibody and a mixture containing an anti-C4 and anti-C4d antibody. Control reactions containing C3b, FI, and FH as well as C4b, FI, and C4BP.
