## Supplementary figure 7 for "CipA mediates complement resistance of *Acinetobacter baumannii* by formation of a Factor I-dependent quadripartite assemblage"

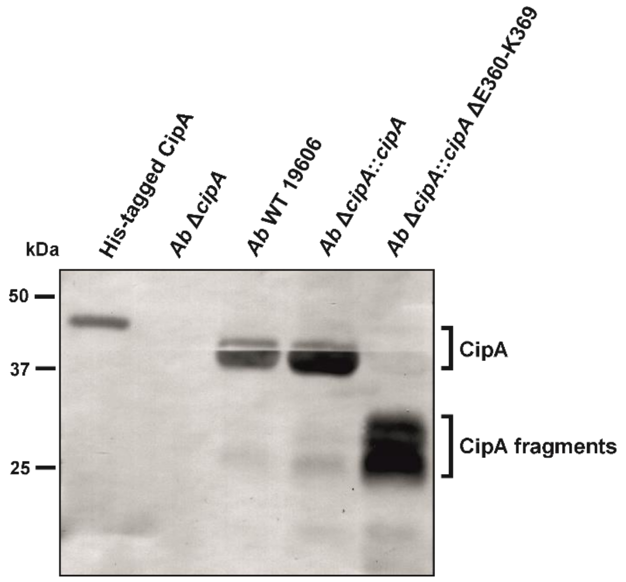


**Fig. S7 Characterization of *A. baumannii* strains producing diverse CipA variants.** CipA was detected in cell lysates (10 µg each) of *A. baumannii* Δ*cipA,* WT 19606 WT, Δ*cipA*::*cipA* and Δ*cipA*::*cipA* ΔE360-K369 by Western blotting using an anti-CipA antibody (1:100). His-tagged CipA was used as control. Native CipA was indicated on the left as well as CipA fragments in strain *A. baumannii* Δ*cipA*::*cipA* ΔE360-K369.
