## Supplementary material for "CipA mediates complement resistance of *Acinetobacter baumannii* by formation of a Factor I-dependent quadripartite assemblage": Legend to supplementary figure 8

**Fig. S8 Structural prediction of CipA and CipA E360P obtained with AlphaFold2.** Structure prediction was created by using the AlphaFold 2 advanced interface with the default settings using the optional “Refine structures with Amber-Relax” option.
