## Supplementary figure 9 for "CipA mediates complement resistance of *Acinetobacter baumannii* by formation of a Factor I-dependent quadripartite assemblage"

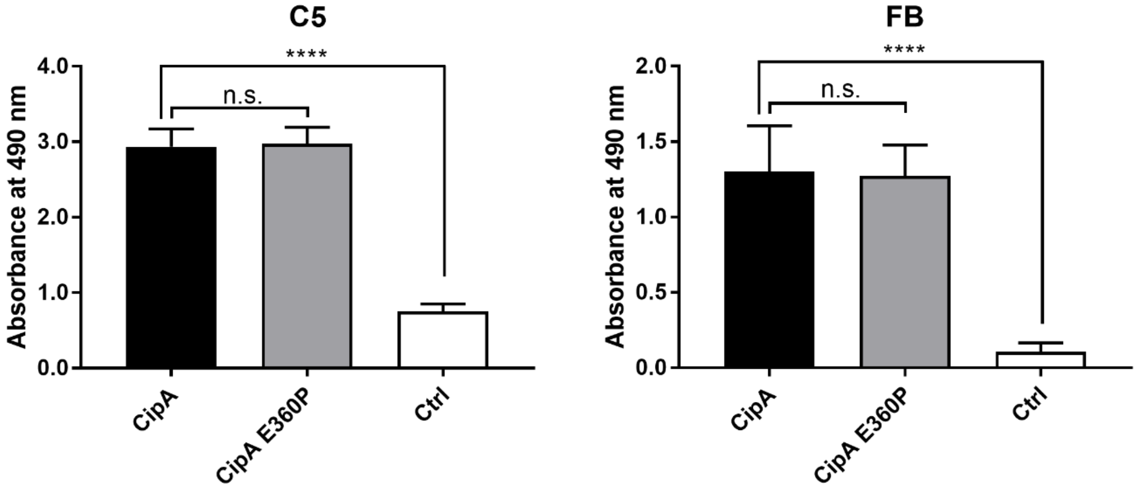


**Fig. S9 Binding of CipA E360P to C5 and FB.** Protein binding of C5 and FB to CipA and CipA E360P was measured by ELISA. Purified bacterial proteins and gelatine (5 ng/µl each) used as negative control were immobilized and incubated with 10 ng/µl C5 or FB. Bound complement components were detected using specific antisera (1:1,000). To assess statistical significance, one-way ANOVA with post-hoc Bonferroni multiple comparison test (confidence interval = 95%) was performed. Data represent means and/or standard deviation of at least three different experiments, each conducted in at least triplicate. ****, $p \leq 0.0001$; n.s., no statistical significance.
