## Supplementary table 1 for "CipA mediates complement resistance of *Acinetobacter baumannii* by formation of a Factor I-dependent quadripartite assemblage"

### Supplementary table 1. Oligonucleotides used in this study

| Oligonucleotide | Sequence (5'-3')[a] | Use in this work |
|---|---|---|
| **Generation of CipA variants with single aa substitutions, deletions in various regions, and extensions** | | |
| CipA-359(-) | TTTTTTAGGCTTTTTGTCGACTTCTTATACAACCATATCTAACACATC | Generation of $CipA_{19-359}$ (aa 19-359), C-terminal deletion of aa 359-369 |
| CipA-145(+) | GCATATTTTCCAAGGATCCAAAGCAACTAATACTCAAGC | Generation of $CipA_{145-369}$ (aa 145-369), N-terminal deletion of aa 19-144 |
| CipA E360A FP | GTGTTAGATATGGTTGTAGCGCAAGAACTCGTTAAAAAAGCC | Generation of variant $CipA_{E360A}$ |
| CipA E360A RP | CTTTTTAACGAGTTCTTGCGCTACAACCATATCTAACACATC | Generation of variant $CipA_{E360A}$ |
| CipA E360P_II FP | GATGTGTTAGATATGGTTGTACCGCAAGAACTCGTTAAAAAGCC | Generation of variant $CipA_{E360P}$ |
| CipA E360P_II RP | CTTTTTAACGAGTTCTTGCGGTACAACCATATCTAACACATCAGC | Generation of variant $CipA_{E360P}$ |
| CipA Q361A FP | GTGTTAGATATGGTTGTAGAGGCAGAACTCGTTAAAAAGCC | Generation of variant $CipA_{Q361A}$ |
| CipA Q361A RP | CTTTTTAACGAGTTCTGCCTCTACAACCATATCTAACACATC | Generation of variant $CipA_{Q361A}$ |
| CipA E362A FP | GATATGGTTGTAGAGCAAGCAACTCGTTAAAAAGCCTAAAAAA | Generation of variant $CipA_{E362A}$ |
| CipA E362A RP | GGCTTTTTAACGAGTGCTTGCTCTACAACCATATCTAACAC | Generation of variant $CipA_{E362A}$ |
| CipA P367A FP | CAAGAACTCGTTAAAAAGGCTAAAAAATAAGTCGACCTGC | Generation of variant $CipA_{P367A}$ |
| CipA P367A RP | GCTTGGCTGCAGGTCGACTTATTTTTTAGCCTTTTTAACGAG | Generation of variant $CipA_{P367A}$ |
| CipA delta L363 FP | GGTTGTAGAGCAAGAAGTTAAAAAGCCTAAAAAATAAGTC | Generation of variant $CipA_{\Delta L363}$ lacking a leucine at aa position 363 |
| CipA delta L363A RP | AGGCTTTTTAACTTCTTGCTCTACAACCATATCTAACAC | Generation of variant $CipA_{\Delta L363}$ lacking a leucine at aa position 363 |
| CipA delta V364 FP | GTAGAGCAAGAACTCAAAAAGCCTAAAAAATAAGTCGACCTG | Generation of variant $CipA_{\Delta V364}$ lacking a valine at aa position 364 |
| CipA delta V364 RP | GAGTTATTTTTTAGGCTTTTTGAGTTCTTGCTCTACAACC | Generation of variant $CipA_{\Delta V364}$ lacking a valine at aa position 364 |
| CipA delta P367 FP | CAAGAACTCTTAAAAAGAAAAAATAAGTCGACCTGC | Generation of variant $CipA_{\Delta P367}$ lacking a proline at aa position 367 |
| CipA delta P367 RP | GCTTGGCTGCAGGTCGACTTATTTTTTCTTTTTAACGAG | Generation of variant $CipA_{\Delta V367}$ lacking a proline at aa position 367 |
| CipA delta L363-V364 FP | GTTGTAGAGCAAGAAAAAAAGCCTAAAAAATAAGTCGACCTG | Generation of variant $CipA_{\Delta L363-V364}$ lacking leucine and valine at positon 363 and 364 |
| CipA delta L363-V364 RP | GAGTTATTTTTTAGGCTTTTTTTCTTGCTCTACAACC | Generation of variant $CipA_{\Delta L363-V364}$ lacking leucine and valine at positon 363 and 364 |
| CipA-364 FP | GTAGAGCAAGAACTCGTTTAATAACCTAAAAAATAAGTCGAC | Generation of variant $CipA_{\Delta V364-K369}$ lacking aa at positions 364 to 369 |
| CipA-364 RP | GTAGAGCAAGAACTCGTTTAATAACCTAAAAAATAAGTCGAC | Generation of variant $CipA_{\Delta V364-K369}$ lacking aa at positions 364 to 369 |
| CipA-L363 FP | GTTGTAGAGCAAGAACTCTAATAAAAGCCTAAAAAATAAGTCGAC | Generation of variant $CipA_{\Delta L363-K369}$ lacking aa at positions 363 to 369 |
| CipA-L363 RP | CGACTTATTTTTTAGGCTTTTATTAGAGTTCTTGCTCTAC | Generation of variant $CipA_{\Delta L363-K369}$ lacking aa at positions 36 to 369 |
| CipA-E362 FP | GTTGTAGAGCAAGAATAATAAAAAAAGCCTAAAAAATAAGTC | Generation of variant $CipA_{\Delta E362-K369}$ lacking aa at positions 362 to 369 |
| CipA-E362 RP | CTTATTTTTTAGGCTTTTTTTATTATTCTTGCTCTAC | Generation of variant $CipA_{\Delta E362-K369}$ lacking aa at positions 362 to 369 |
| CipA-Q361 FP | GGTTGTAGAGCAATAATAAGTTAAAAAGCCTAAAAAAT | Generation of variant $CipA_{\Delta Q361-K369}$ lacking aa at positions 361 to 369 |

| | | |
|---|---|---|
| CipA-Q361 RP | GGCTTTTTAACTTATTATTCCTCTACAACCATATC | Generation of variant CipA$_{\Delta Q361\text{-}K369}$ lacking aa at positions 361 to 369 |
| CipA-E360 FP | GATATGGTTGTAGAGTAATAACTCGTTAAAAAGCCTAAAAAAT | Generation of variant CipA$_{\Delta E360\text{-}K369}$ lacking aa at positions 360 to 369 |
| CipA-E360 RP | GGCTTTTTAACGAGTTATTACTCTACAACCATATCTAAC | Generation of variant CipA$_{\Delta E360\text{-}K369}$ lacking aa at positions 360 to 369 |
| **Generation of *A. baumannii* 19606 strain producing a CipA variant by markerless mutagenesis** | | |
| CipA up fwd PstI | GCGACTGCAGCAAACTCAGGTTATTGAACTCCCAATGG | Amplification of the upstream region of the CipA encoding gene in strain *A. baumannii* 19606 |
| CipA down rev NotI | GACAGCGGCCGCGGGATTTAATATTTTGCTGCTAAATG | Amplification of the downstream CipA encoding gene in strain *A. baumannii* 19606 |
| CipA V359 RP | GTTAGATATGGTTGTATAGTAATAACTCGTTAAAAAGCCTAAAAAAA | Introduction of two stop codons after position 359 |
| CipA V359 FP | GGCTTTTTAACGAGTTATTACTATACAACCATATCTAACACATC | Introduction of two stop codons after position 359 |
| CipA control fwd | GCTCTTGTCTATCTTATGTCACAGATAGCC | Control of the complementation in *A. baumannii* 19606 |
| CipA control rev | CTGGCTGTCCACCAGGAACACATTTATTTGC | Control of the complementation in *A. baumannii* 19606 |
| CipA seq Fwd | CAATCCAAACCAACGCGTAATCTTACGTG | Sequencing of *cipA* gene in the complemented *A. baumannii* 19606 |
| CipA seq Rev | GGTACTGTGAATTATGTAGATGGTGCC | Sequencing of *cipA* gene in the complemented *A. baumannii* 19606 |
| **Primer used for RT-PCR of *A. baumannii* genes** | | |
| *rpoB*-RT-fwd | GAGTCTAATGGCGGTGGTTC | RT-PCR, amplification of *rpoB* in *A. baumannii* |
| *rpoB*-RT-rev | ATTGCTTCATCTGCTGGTTG | RT-PCR, amplification of *rpoB* in *A. baumannii* |
| CipA-RT-PCR Fwd | GTCATGTCAACTATGATGGC | RT-PCR, amplification of *cipA* in *A. baumannii* |
| CipA-RT-PCR Rev | GCAAGCAAGTTTGTGGTGCAAC | RT-PCR, amplification of *cipA* in *A. baumannii* |
| **Primer used for the generation of His-tagged Efb and Efb-C proteins of *Staphylococcus aureus* USA300** | | |
| Efb_FP Bam | GCGAGCGAAGGATCCGGTCCAAGAGAAAAGAAACCAGTGAG | Amplification of Efb encoding gene of *S. aureus* USA300 |
| Efb_RP_Sal | GGCTGCAGGTCGACTTATTTAACTAATCCTTGTTTTAATAC | Amplification of Efb encoding gene of *S. aureus* USA300 |
| Efb-C_FP_Bam | GGTGCAGGATCCCAATTTAATAAACCAGCAGCGAAAACTG | Generation of a C-terminal fragment of Efb |
| **Primer used for sequencing** | | |
| pQE-FP-30 | TTGCTTTGTGAGCGGATAAC | Sequencing of pQE-30 Xa vector |
| pQE-RP | CTGAGGTCATTACTGGATCTATC | Sequencing of pQE-30 Xa vector |

[a], Sequences of specific restriction endonuclease recognition sites are underlined
